# Polyploidization events shaped the transcription factor repertoires in legumes (Fabaceae)

**DOI:** 10.1101/849778

**Authors:** Kanhu C. Moharana, Thiago M. Venancio

**Author notes:** Corresponding author Av. Alberto Lamego 2000 / P5 / 217; Parque Califórnia Campos dos Goytacazes, RJ Brazil CEP: 28013-602. ***Abbreviations*** **TF:** transcription factors; **DBD:** DNA binding domains; **AP2:** APETALA 2; **ERF:** Ethylene Responsive Factor; **RAV:** Related to ABI3/VP1; **ARF:** Auxin response factor; **BBR-BPC:** Barley B Recombinant (BBR) - BASIC PENTACYSTEINE1 (BPC1); **BES1:** BRI1-EMS-SUPPRESSOR; **bHLH:** Basic helix loop helix; **bZIP:** Basic leucine zipper; **Dof:** DNA binding with one finger; **CO-like:** CONSTANS-like; **LSD:** LESION SIMULATING DISEASE 1 (LSD1); **C2H2:** CCHH (Zn); **C3H:** CCCH (Zn); **CAMTA:** Calmodulin binding transcription factors; **CPP:** Cystein-rich polycomb-like protein; **DBB:** Double B-box zinc finger; **E2F/DP:** E2 factor protein and DP protein; **EIL:** Ethylene-Insentive3 (EIN3)-like protein3 (EIL3); **FAR1:** FAR-RED IMPAIRED RESPONSE1; **LFY:** LEAFY; **G2-like:** Golden2 (G2)-like; **ARR-B:** Type-B phospho-accepting response regulator; **GeBP:** GLABROUS1 enhancer-binding protein; **GRAS:** GAI, RGA, and SCR; **GRF:** GROWTH-REGULATING FACTOR; **TALE:** Three Amino acid Loop Extension; **WOX:** WUS homeobox-containing protein family; **HB:** Homeobox **; HB-PHD:** HB-PHD finger; **HB-other:** HB-other; **HRT-like:** Hairy-Related transcription-factor-like; **HSF:** Heat shock factor; **LBD:** ASYMMETRIC LEAVES2/LATERAL ORGAN BOUNDARIES; **M-type:** MADS-type I; **MIKC:** MADS-type II; **MYB:** Myb proto-oncogene protein; **NAC:** NAM, ATAF1, 2 and CUC2; **NF-X1:** Nuclear factor, X-box binding 1; **NF-YA:** Nuclear factor Y subunit A; **NF-YB:** Nuclear factor Y subunit B; **NF-YC:** Nuclear factor Y subunit C; **Nin-like:** NODULE INCEPTION; **NZZ/SPL:** SPOROCYTELESS/NOZZLE; **S1Fa-like:** S1Fa-like; **TCP:** TEOSINTE-LIKE1, CYCLOIDEA, and PROLIFERATING CELL FACTOR1; **ZF-HD:** Zinc finger homeodomain protein; **SBP:** SQUAMOSA promoter binding protein; **SRS:** SHI RELATED SEQUENCE; **SAP:** STERILE APETALA; **STAT:** Signal Transducers and Activators of Transcription; **VOZ:** Vascular plant One-Zinc finger.

## Abstract

Transcription factors (TF) are essential for proper plant growth and development. Several legumes, particularly soybean, are rich sources of protein and oil, with great impact in the economy of several countries. Here we report a phylogenomic analysis of major TF families in legumes and their potential association with important traits such as nitrogen fixation and seed development. We used TF DNA-binding domains to systematically screen the genomes of 15 legume and 5 non-legume species. The percentage of TFs ranged from 3-8% of the gene complements. TF orthologous groups (OG) in extant species were used to estimate OG sizes in ancestor nodes using a gene birth-death model, which allowed us to identify lineage-specific expansions. Together, OG analysis and rate of synonymous substitutions (Ks) between gene pairs show that major TF expansions are strongly associated with known whole-genome duplication (WGD) events in the legume (∼58 mya) and Glycine (∼13 mya) lineages, which account for a large fraction of the *Ph. vulgaris* and *Gl. max* TF repertoires. Out of the 3407 *Gl. max* TFs, 1808 and 676 can be traced back to a single homeolog in *Ph. vulgaris* and *Vi vinifera*, respectively. We found a trend for TFs expanded in legumes to be preferentially transcribed in roots and nodules, suggesting their recruitment early in the evolution of nodulation in the legume clade. We also found TF expansions in the Glycine WGD that were followed by gene loss in the wild soybean *Gl. soja,* including genes located within important quantitative trait loci. Together, our findings strongly support the roles of two WGDs in shaping the TF repertoires in the legume and Glycine lineages, which are likely related to important aspects of legume and soybean biology.

## Introduction

Legumes (Fabaceae) are the third largest Angiosperm family, comprising nearly 20,000 species with tremendous morphological and ecological variation (Lewis, 2005). Legumes are notorious for their symbiotic interactions with specific diazotrophic bacteria, which is a feature of major ecological and agronomic relevance. Out of the six Fabaceae subfamilies, Papilionoideae alone accounts for two-thirds of all legume species, including economically important crops, such as *Glycine max* (soybean), *Phaseolus vulgaris* (common bean), *Arachis hypogaea* (peanut), and *Cicer arietinum* (chickpea) (Graham and Vance, 2003;Cardoso et al., 2012;Azani et al., 2017). Several legume grains and pulses are rich sources of dietary proteins, cooking oils, and biofuels. Currently, at least 15 legume genomes are publicly available (Table 1), including those from wild and cultivated soybean (*Glycine soja* and *Gl. max*). Nevertheless, Fabaceae subfamilies other than Papilionoideae are largely underrepresented among sequenced genomes, despite the recently published *Chamaecrista fasciculata* and *Mimosa pudica* genomes (Griesmann et al., 2018).

**Table 1:** Plant species used in this study and their corresponding genome assembly versions. Non-legume species are marked with asterisks.

| Scientific name | Genome assembly version | Genes | Chromosome number | Reference |
| --- | --- | --- | --- | --- |
| *Cajanus cajan* | GCA 000340665.1 | 48,331 | 11 | (Varshney et al., 2011) |
| *Phaseolus vulgaris* | Pvulgaris 218 v1.0 | 27,197 | 11 | (Schmutz et al., 2014) |
| *Vigna radiata* | Vradiata ver6 | 35,143 | 11 | (Kang et al., 2014) |
| *Vigna angularis* | Vigan1.1 | 34,172 | 11 | (Yang et al., 2015) |
| *Glycine max* | Gmax 275 Wm82.a2.v1 | 56,044 | 20 | (Schmutz et al., 2010) |
| *Glycine soja* | W05v1.0 | 55,539 | 20 | (Xie et al., 2019) |
| *Cicer reticulatum* | WCGAP v1.0 | 25,680 | 8 | (Gupta et al., 2017) |
| *Cicer arietinum* | ASM33114v1 | 33,107 | 8 | (Jain et al., 2013;Varshney et al., 2013;Parween et al., 2015) |
| *Medicago truncatula* | Mtruncatula 285 Mt4.0v1 | 50,894 | 8 | (Young et al., 2011;Tang et al., 2014) |
| *Glycyrrhiza uralensis* | Gur.draft-genome.20151208 | 34,445 | 8 | (Mochida et al., 2017) |
| *Lotus japonicus* | Genome assembly build 3.0 | 39,734 | 6 | (Sato et al., 2008) |
| *Lupinus angustifolius* | v1.0 | 33,076 | 20 | (Hane et al., 2017) |
| *Arachis ipaensis* | Araip1.0 | 46,410 | 10 | (Bertioli et al., 2016) |
| *Arachis duranensis* | Aradu1.0 | 42,562 | 10 | (Bertioli et al., 2016;Chen et al., 2016) |
| *Chamaecrista fasciculata* | version.1 | 21,781 | 8 | (Griesmann et al., 2018) |
| *Arabidopsis thaliana* * | Athaliana 167 TAIR10 | 27,416 | 5 | (Arabidopsis-Genome-Initiative, 2000) |
| *Vitis vinifera* * | Vvinifera 145 Genoscope.12X | 26,346 | 19 | (Jaillon et al., 2007) |
| *Amborella trichopoda* * | AmTr v1.1 | 26,846 | 13 | (Albert et al., 2013) |
| *Aquilegia coerulea* * | Aquilegia coerulea v3.1 | 30,023 | 7 | (Filiault et al., 2018) |
| *Selaginella moellendorffii* * | Selaginella moellendorffii v1.0 | 22,285 | 10 | (Banks et al., 2011) |

Virtually all major biological processes are at least partially regulated at the transcriptional level by specific DNA-binding transcription factors (TFs), which bind to *cis*-regulatory elements of target genes by means of DNA binding domains (DBDs). Because of their key regulatory roles, TFs have been extensively demonstrated to be critical for plant evolution and adaptation to multiple environments (Doebley and Lukens, 1998;Lehti-Shiu et al., 2017). A typical TF family encodes proteins sharing a common DBD. Over 50 TF families have been identified in plants (Jin et al., 2017), out of which many regulate biological processes such as growth, development, stress signaling, and defense against pathogens (Lata et al., 2011;Tripathi et al., 2016;He et al., 2018). Although many TF families are present throughout eukaryotes, their sizes can vary considerably (Lespinet et al., 2002;Nagata et al., 2016;Lehti-Shiu et al., 2017). Plant TF families are usually larger than their animal counterparts (Shiu, 2005) and the exceptional increase in TF family sizes in higher plants is often related with whole genome duplication (WGD) and triplication (WGT) events, also known as polyploidization (Lehti-Shiu et al., 2017).

Polyploidization is widely accepted as an important factor in angiosperm evolution (Pickett and Meeks-Wagner, 1995;Soltis et al., 2015). All core eudicots share at least one polyploidization event (a hexaploidy event, i.e. the γ polyploidy) (Jaillon et al., 2007;Aköz and Nordborg, 2019). With the increasing availability of plant genomes, it became clear that several other WGD events occurred after the γ polyploidization event. For example, *Arabidopsis* has two additional lineage-specific WGDs (α and β polyploidy) (Blanc et al., 2000;Vision et al., 2000), whereas legumes share a tetraploid ancestor that originated ∼58-60 million years ago (mya) (Cannon et al., 2006). Further analysis of the soybean and narrowleaf lupin (*Lupinus angustifolius*) genomes uncovered additional lineage-specific WGDs in these lineages (Schmutz et al., 2010;Kroc et al., 2014;Hane et al., 2017). Importantly, domestication of polyploid species is more likely than that of their wild relatives (Salman-Minkov et al., 2016), supporting the importance of such events in agriculture. In addition to large scale duplication events, small scale duplications (SSDs) or local (e.g. tandem) duplications also contributed to the expansion of multiple gene families (Cannon et al., 2004).

The relative contribution of duplication modes in plant genomes is a subject of intense research (Panchy et al., 2016). Upon WGD, gene loss is the most common fate of one of the duplicates, in a process called fractionation (Freeling et al., 2015;Panchy et al., 2016;Cheng et al., 2018). Nevertheless, some gene families (e.g. TFs and signal transduction genes) are apparently more prone to retain duplicated copies than others (Blanc and Wolfe, 2004;Lehti-Shiu et al., 2017). Different mechanistic explanations have been proposed for this phenomenon, out of which the *gene balance hypothesis* is the most accepted one. According to this hypothesis, upon a WGD, genes with many interaction partners have higher probability of being retained in duplicates, since alterations in the stoichiometry their protein products tend to be deleterious (Birchler and Veitia, 2007;Freeling, 2009;Birchler and Veitia, 2011). Retained copies then typically evolve via subfunctionalization (i.e. duplicates acquire complementary functions) or neofunctionalization (i.e. one of the copies evolves a new function) (Freeling et al., 2015).

It is currently accepted that the transition from wild to domesticated soybean took place in Central China between 5,000 and 9,000 years ago through a gradual process that involved an intermediary species, *Gl. gracilis* (Han et al., 2016;Sedivy et al., 2017). Other lines of evidence support independent domestication events in East Asia (Korea and Japan) (Zhou et al., 2015;Sedivy et al., 2017). Artificial selection during domestication involved several distinct traits, such as pod shattering, seed hardness, adaptation to different photoperiods, flowering time, and stress resistance (Zhou et al., 2015;Sedivy et al., 2017). Several important TFs are involved in these traits. *SHAT1-5* (NAC family TF) promote shattering resistance by increasing lignification of fiber cap cells (Dong et al., 2014).

The progress in sequencing technologies and automation over the past 12 years unleashed the power of comparative and population genomics in pinpointing key genes involved in domestication and improvement of soybean and other crops, as illustrated by the discovery of many Quantitative Trait Loci (QTL) involved in commercially important traits (e.g. seed weight and oil content) by the resequencing of 302 soybean accessions (Zhou et al., 2015). In the present work we systematically investigate the evolution of TF repertoires in legumes (Fabaceae). Briefly, we performed large-scale comparative analysis of TFs from 15 legume and five non-legume species. Our results unveil a profound impact of polyploidization events on the expansion of TF families throughout legumes. In particular, some TF families that expanded in the legume WGD event (∼58 mya) are preferentially expressed in roots and nodules, supporting their importance in the evolution of nodulation. Further, TF expansions that happened at the Glycine WGD (∼13 mya) include genes that were subsequently lost in *Gl. soja*, including TF genes that are within *Gl. max* QTLs associated with leaf shape, area and width, proportion of FA18 in seeds, and branch density. Together, our results strongly support that WGD events deeply shaped the evolution of TF repertoires in legumes and likely generated TFs that regulate nodulation and other traits of key agronomic importance.

## Results and Discussion

### Systematic identification and characterization of transcription factors

We used a set of diagnostic specific DNA binding domains and forbidden domains (Supplementary Table S1) to identify TFs in the genomes of 20 plant species (Table 1). We predicted a total of 37,008 TFs (Supplementary Table S2), which were classified in 58 broad families (Table 2). A total of 31,111 TFs were predicted in the 15 legume genomes (Supplementary Table S2). We benchmarked our pipeline by comparing the detected TFs with those previously predicted in *Ar. thaliana*. Out of 1713 *Ar. thaliana* TFs available in PlantTFDB, 1673 (98%) were correctly predicted (Supplementary Figure S1). Further, 59 TFs were exclusively predicted by our pipeline, out of which 40 were annotated as TFs in the TAIR database (https://www.arabidopsis.org) (Supplementary Table S3). The percentage of TFs across genomes ranged from 5 to 8%, which is in line with a previous estimation from 95 eudicot species (Jin et al., 2017). *Gl. soja* and *Se. moellendorffii* showed the highest and lowest number of TFs, respectively (Figure 1). Legumes typically showed greater number of TFs than non-legumes (Figure 1), although the variation in these fractions indicates that some TF expansions play specific roles in particular lineages.

**Figure 1:**
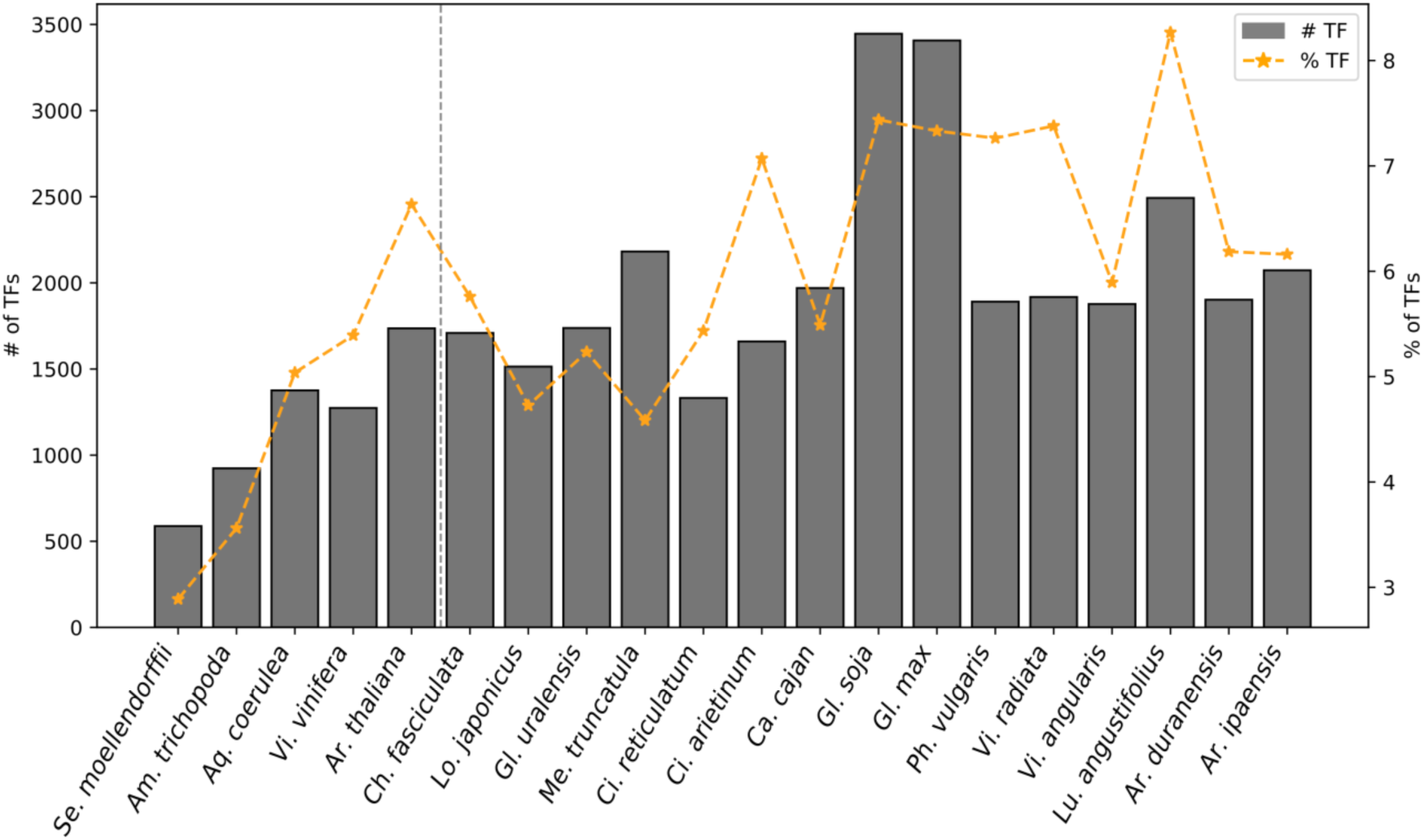
Absolute and relative number of transcription factors in each species. Grey bars and the orange line represent the absolute number and percentage of transcription factors in each species, respectively. Legumes and non-legumes are separated by a dotted vertical line.

**Table 2:**
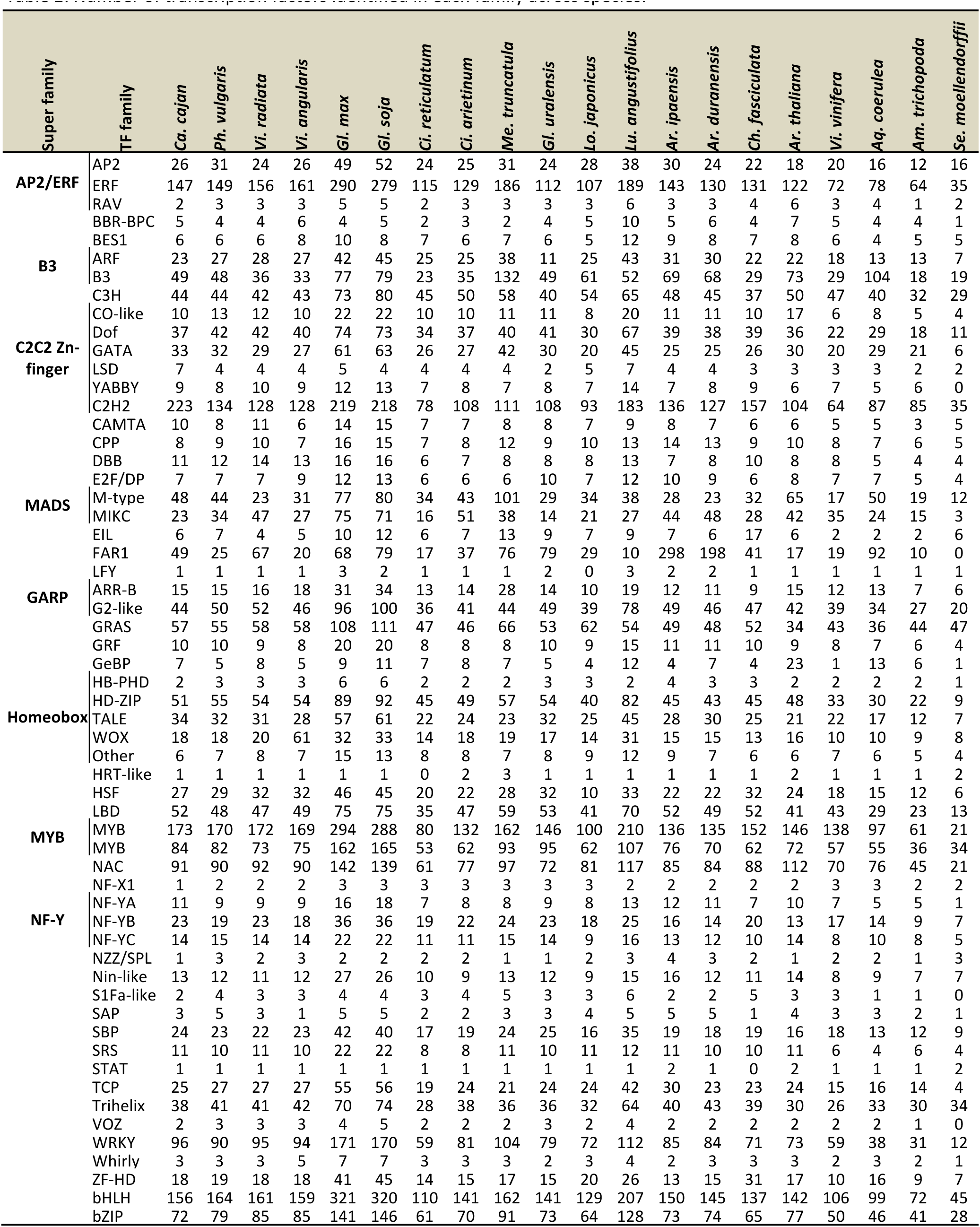
Number of transcription factors identified in each family across species.

To better understand the different proportion of TFs across genomes, we compared TF family sizes between pairs of species. All except six TF families (i.e. AP2, GRAS, B3, Nin-like, HRT-like, and Trihelix) expanded in the basal angiosperm *Am. trichopoda* in comparison to the lycophyte *Se. moellendorffii*, supporting the contribution ancient WGD events to the TF repertoires of seed plants (Albert et al., 2013). Although MADS TFs tightly regulate flower development, their diversification has been proposed to predate the origin of angiosperms (Albert et al., 2013). We found twice more MADS genes in *Am. trichopoda* (n=34) than in *Se. moellendorffii* (n=15). In particular, the MIKC-type (type II) MADS subfamily alone has increased by five-fold, in spite of the higher rate of gene birth/death of the M-type (type-I) MADS subfamily (Nam et al., 2004;Kumpeangkeaw et al., 2019). By analyzing TF clusters (described later) we observed that genes from two M-type MADS clusters are exclusively present in *Am. trichopoda*, probably as a result of a lineage-specific expansion. We also analyzed the expansion of the GRAS family in *Am. trichopoda* (n= 44) as compared to the basal dicot *Aq. coerulea* (n= 36), which happened via lineage-specific tandem duplications (10 genes) in the former (Supplementary Figure S3). These 10 genes belong to a single orthologous group (OG, see below) that does not have orthologs from other dicots except one from the basal eudicot *Aq. coerulea*. In addition, we found one more *Am. trichopoda* specific OG consisting of two GRAS genes (scaffold00007.332 and scaffold00007.335). There are also a few remarkable expansions in *Se. moellendorffii* in comparison to *Am. trichopoda* (e.g. HD-Zip, NAC, TCP, GATA, expanded by more than two fold) (Table 2). Together, these results support a growth of the TF repertoire early in the diversification of angiosperms.

*Aq. coerulea* is an ancient tetraploid and this tetraploidy was likely an important first step towards the *gamma* hexaploidy (4n+2n) that is shared by all core eudicots (Aköz and Nordborg, 2019). Nevertheless, some TF families are remarkably larger in *Aq. coerulea* than in *Vi. vinifera*, such as FAR1 (Aco: 92, Vvi: 19), B3 (Aco: 104, Vvi:29), GeBP (Aco: 13, Vvi: 1), and M-Type MADS (Aco: 50, Vvi: 17) (Figure 2; Table 2). After the *gamma* hexaplodization event, *Vi. vinifera* has not undergone any large scale duplication event, making it a suitable reference for comparative analysis with other core eudicots (Jaillon et al., 2007;Severin et al., 2011;Wang et al., 2017). Most large families are expanded in *Ar. thaliana* and legumes in comparison to *Vi. vinifera* (Figure 2). There are also some notable species-specific expansions in legumes, such as that of FAR1, B3, and M-Type MADS in *Me. truncatula* (Figure 2). FAR1 has also expanded 10 times in *Ar. ipaensis* and *Ar. duranensis*. FAR1 has been linked with skotomorphogenesis and photomorphogenesis in higher plants and its expansion might be related with the fructification process in peanuts (Chen et al., 2016;Lu et al., 2018). Unlike the above-mentioned expansions of MADS TFs in *Am. trichopoda,* only M-type (type-I) MADS had large expansions in all legumes in comparison to *Vi. Vinifera*, as previously discussed (Nam et al., 2004;Feil et al., 2013;Kumpeangkeaw et al., 2019).

**Figure 2:**
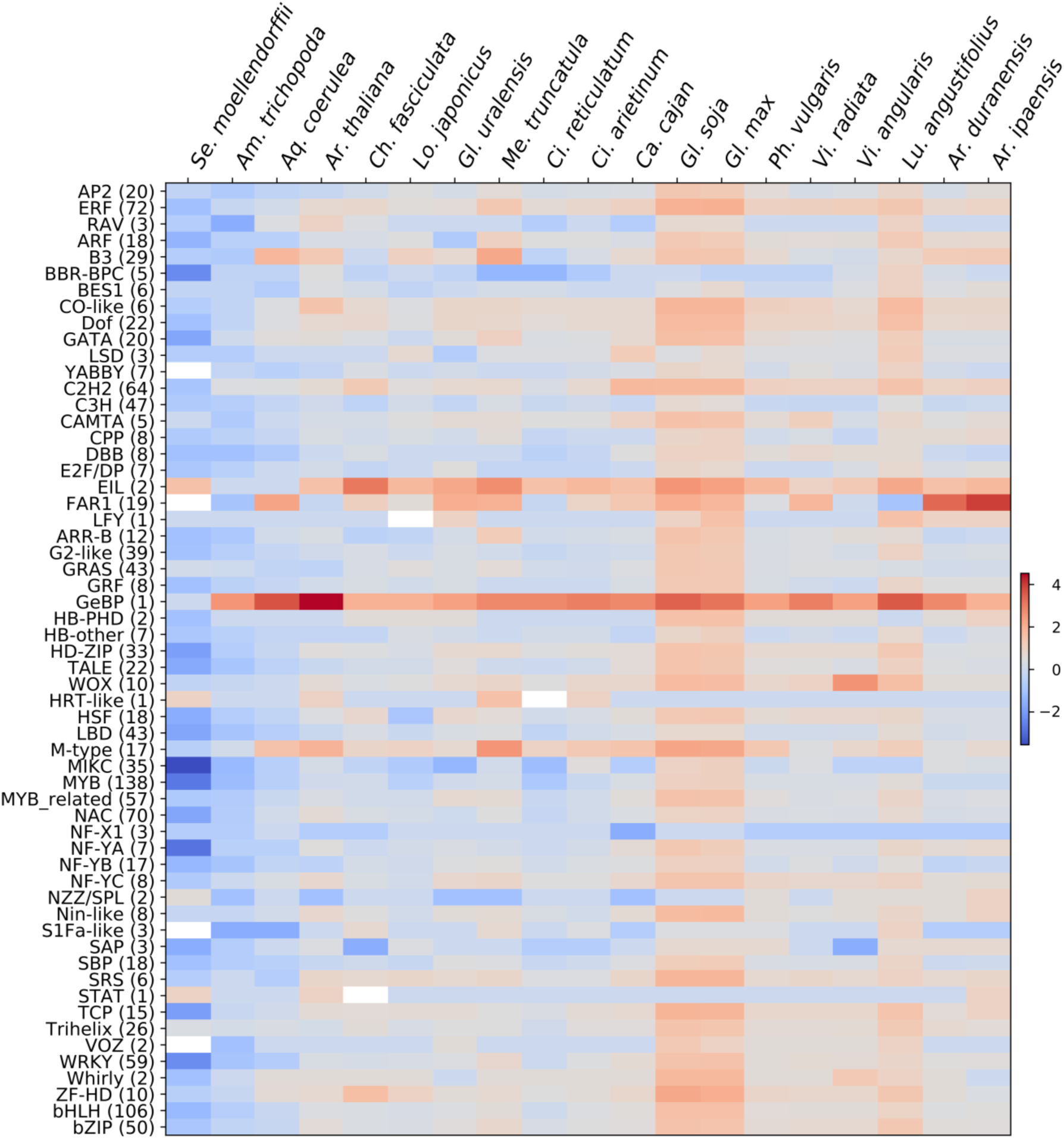
Ratio (in log_2_ scale) of the sizes of each transcription factor family each species in relation to *Vi. vinifera*. Values greater or smaller than zero represent transcription factor families that are relatively larger or smaller in a given species (in columns) in comparison to *Vi. vinifera*, respectively. The numbers in parentheses stand for the absolute size of that particular family in *Vi. vinifera*.

The *Glycine* genus has a more recent WGD that is not shared with *Phaseolus*. Accordingly, we found an approximate ratio of 1:2 between *Ph. vulgaris* and *Gl. max* in 90% (52/58) of the TF families, implying that the *Glycine* WGD has strongly contributed to the soybean TF repertoire. There are also a few deviations from this trend, such as the NAC (*Gl. max*: 142 and *Ph. vulgaris*: 90) and NOZZLE/SPL (*Gl. max*: 2 and *Ph. vulgaris:* 3) families. Of the 42 NAC OGs with genes from *Ph. vulgaris* and *Gl. max*, 9 have identical number of genes, indicating that there are subfamilies that rapidly reverted back to their configuration before the *Glycine* WGD, probably due to gene dosage sensitivity. We also noticed that *Gl. soja* has 38 more TFs than *Gl. max*. While sixteen TF families had identical number of genes in both species, others such as ERF (*Gl. max*: 290, *Gl. soja*: 279) and FAR1 (*Gl. max*: 68, *Gl. soja*:79) showed significant variation between cultivated and wild soybeans.

### Origin of TF paralogs

We also investigated the modes of duplication shaping TF family sizes. Paralogous pairs had their modes of duplication predicted using a previously described scheme (Proulx et al., 2011;Qiao et al., 2018) that assign duplicate pairs in the following categories: segmental duplicates (SD); tandem duplicates (TD); pairs separated by one to five intervening genes were called proximal duplicates (PD); pairs originated via retrotransposon (rTE) or DNA transposon (dTE) activity. The remaining pairs were classified as dispersed duplicates (DD). We used the priority order SD>TD>PD>rTE>dTE>DD to assign a single duplication mode to each pair. In legumes, more than 70% of the TFs have at least one paralog (Table 3). Further, it is clear that SD is the main duplication mode, supporting their origin through large scale duplication, as previously reported (Lehti-Shiu et al., 2017). *Gl. max* and *Gl. soja* are the species with the greatest number of SD TFs, which comprise 77.7% of the TF repertoire in the former. In addition, local duplications (i.e. TD and PD) also contributed to TF repertoires, particularly in *Me. truncatula*, which has 11% (241/1752) of the duplicate TFs classified as TD and PD, especially in the ERF, WRKY, and B3 families (Supplementary Table S4). The prevalence of TD pairs in *Me. truncatula* has also been reported in genes related to other regulatory roles, such as in the F-box family (Bellieny-Rabelo et al., 2013). Further, in *Ar. duranensis, Ar. ipaensis*, and *Vi. radiata*, nearly 7% of the duplicated TFs are derived from TD, whereas dTE duplications account for 29.8% of the duplicated TFs in *Ca. fasciculata*, especially in the MYB, NAC, and bHLH families (Table 3; Supplementary Table S4). There are also some notable differences in the prevalence of modes of duplication between closely related species. For example, local TF duplications are more frequent in *Ar. ipaensis* than in *Ar. duranensis* (Table 3).

**Table 3:** Prevalence of different modes of duplication among transcription factors.

| Species | Total | Singletons (%) | Duplicates | SD | TD | PD | rTE | dTE | DD |
| --- | --- | --- | --- | --- | --- | --- | --- | --- | --- |
| *Ca. cajan* | 1970 | 314 (15.94) | 1656 (84.06) | 548 | 90 | 21 | 40 | 397 | 560 |
| *Ph. vulgaris* | 1891 | 257 (13.59) | 1634 (86.41) | 914 | 99 | 8 | 16 | 244 | 353 |
| *Vi. radiata* | 1918 | 253 (13.19) | 1665 (86.81) | 743 | 115 | 9 | 24 | 319 | 455 |
| *Vi. angularis* | 1877 | 307 (16.36) | 1570 (83.64) | 842 | 74 | 9 | 14 | 220 | 411 |
| *Gl. max* | 3407 | 188 (5.52) | 3219 (94.48) | 264 | 79 | 14 | 28 | 242 | 211 |
| *Gl. soja* | 3445 | 224 (6.5) | 3221 (93.5) | 264 | 85 | 13 | 24 | 231 | 220 |
| *Ci. reticulatum* | 1332 | 365 (27.4) | 967 (72.6) | 350 | 45 | 3 | 24 | 207 | 338 |
| *Ci. arietinum* | 1660 | 339 (20.42) | 1321 (79.58) | 521 | 101 | 7 | 17 | 267 | 408 |
| *Me. truncatula* | 2182 | 430 (19.71) | 1752 (80.29) | 648 | 209 | 32 | 27 | 235 | 601 |
| *Gl. uralensis* | 1738 | 403 (23.19) | 1335 (76.81) | 482 | 36 | 3 | 19 | 281 | 514 |
| *Lo. japonicus* | 1514 | 428 (28.27) | 1086 (71.73) | 195 | 39 | 6 | 25 | 345 | 476 |
| *Lu. angustifolius* | 2493 | 223 (8.95) | 2270 (91.05) | 165 | 51 | 0 | 0 | 0 | 565 |
| *Ar. ipaensis* | 2073 | 365 (17.61) | 1708 (82.39) | 336 | 113 | 21 | 32 | 434 | 772 |
| *Ar. duranensis* | 1902 | 363 (19.09) | 1539 (80.91) | 410 | 95 | 5 | 27 | 373 | 629 |
| *Ch. fasciculata* | 1709 | 426 (24.93) | 1283 (75.07) | 186 | 36 | 2 | 36 | 383 | 640 |
| *Ar. thaliana* | 1736 | 392 (22.58) | 1344 (77.42) | 707 | 81 | 13 | 15 | 152 | 376 |
| *Vi. vinifera* | 1274 | 478 (37.52) | 796 (62.48) | 303 | 82 | 13 | 5 | 193 | 200 |
| *Aq. coerulea* | 1376 | 552 (40.12) | 824 (59.88) | 130 | 86 | 14 | 0 | 0 | 594 |
| *Am. trichopoda* | 923 | 560 (60.67) | 363 (39.33) | 14 | 33 | 4 | 0 | 0 | 312 |
| *Se. moellendorffii* | 588 | 278 (47.28) | 310 (52.72) | 79 | 10 | 4 | 0 | 0 | 217 |

Abbreviations: Segmental duplicates (SD); Tandem duplicates (TD); Proximal duplicates (PD); Retrotransposon mediated duplicates (rTE); Transposon mediated duplicates (dTE); Dispersed duplicates (DD).

Between 5.5% (188/3407, in *Gl. max*) and 60.7% (560/923, in *Am. trichopoda*) of the TFs were classified as singletons (Table 3). While in *Ar. thaliana* 22.58% (392/1736) of the TFs were singletons, in legumes this number ranges between 5.5 and 29% (Table 3). Importantly, a large fraction of these singletons remain syntenic to a reference outgroup species (Table 4). In *Ph. vulgaris* and *Me. truncatula*, syntenic singleton TFs were significantly more expressed than their non-syntenic counterparts (Figure 3A), suggesting that their greater functional conservation is associated with their genomic context. Most SD TFs were also found to be syntenic in their closest outgroup species (Table 4). We also estimated non-synonymous/synonymous mutation ratios (Ka/Ks) between singleton and SD TFs with preserved synteny in an outgroup species. Orthologs from SD pairs had significantly lower Ka/Ks than the singleton orthologs (Figure 3B), leading us to hypothesize that these genes are under strong purifying selection due to their involvement in intricate regulatory systems emerging from the WGD events. Similar observations on the strong negative selection of duplicated genes have been also reported in other species (Davis and Petrov, 2004;Jordan et al., 2004).

**Figure 3:**
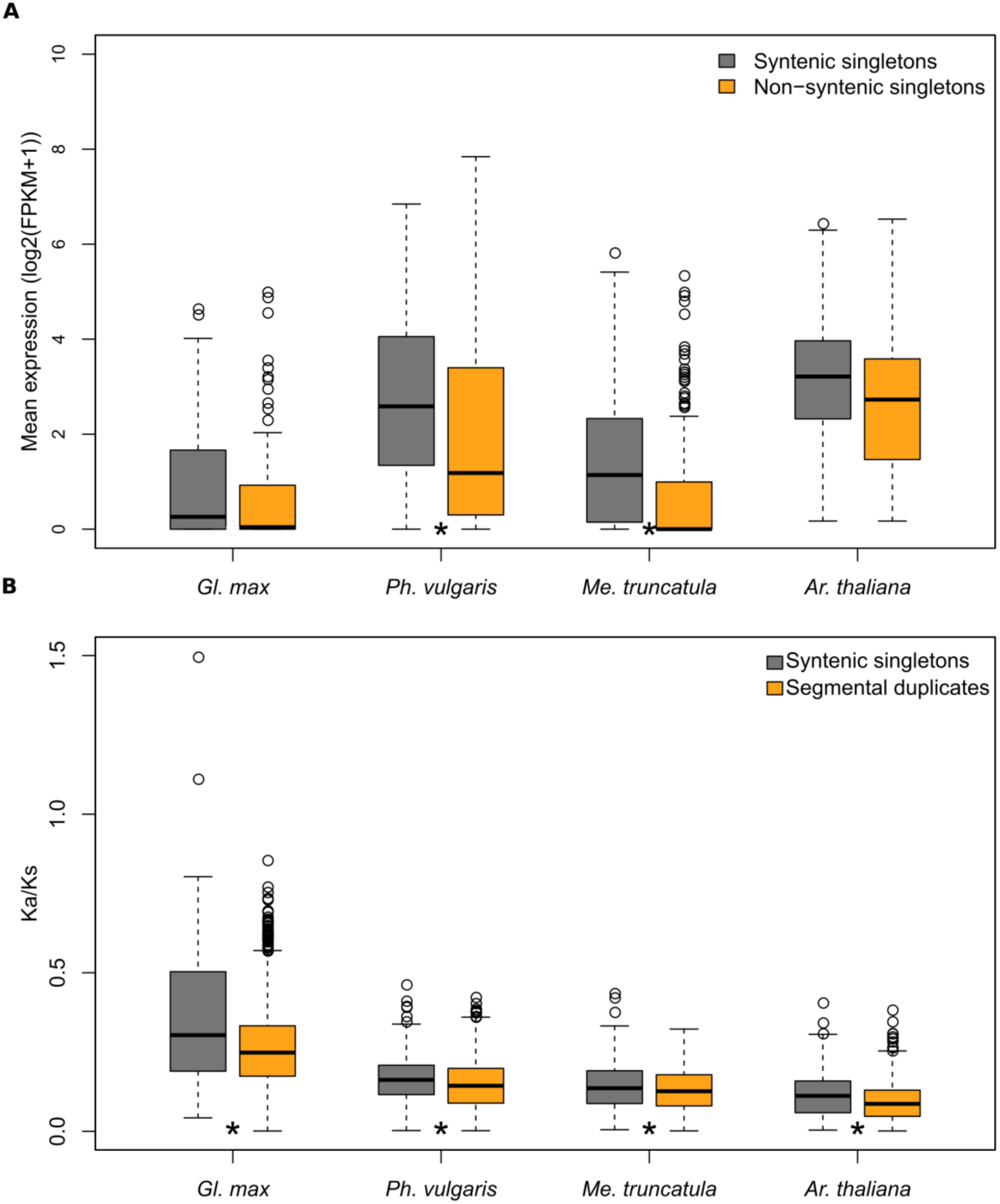
Expression levels and Ka/Ks ratio of singleton transcription factors. A. Expression (in FPKM) of syntenic- and non-syntenic singletons in three legume species and in *Arabidopsis thaliana*. B. Ka/Ks distribution of syntenic singletons and syntenic segmental duplicates. Syntenic singletons are transcription factors genes, without a close paralog, that are located in a syntenic region in a reference outgroup. Segmental duplicates are paralogous transcription factors with preserved synteny in the same genome, as well as in the genome of a reference outgroup. *Ph. vulgaris* was used as reference for *Gl. max* and *Vi. vinifera* was used as reference for the other three species. Statistical significance test was performed using the Mann-Whitney U test and asterisk (*) mark indicates p-value < 0.05.

**Table 4:** Percentage of singletons within collinear regions in a reference species.

| Species | Reference outgroup species | Singletons | Singletons in collinear regions | SD | SD in collinear regions in the reference outgroup |
| --- | --- | --- | --- | --- | --- |
| *Ca. cajan* | *Vi. vinifera* | 314 | 132 (42%) | 548 | 430 (78%) |
| *Ph. vulgaris* | *Vi. vinifera* | 257 | 142 (55%) | 914 | 750 (82%) |
| *Vi. radiata* | *Vi. vinifera* | 253 | 120 (47%) | 743 | 595 (80%) |
| *Vi. angularis* | *Vi. vinifera* | 307 | 161 (52%) | 842 | 659 (78%) |
| *Gl. max* | *Ph. vulgaris* | 188 | 93 (49%) | 2645 | 2546 (96%) |
| *Gl. soja* | *Ph. vulgaris* | 224 | 105 (47%) | 2648 | 2546 (96%) |
| *Ci. reticulatum* | *Vi. vinifera* | 365 | 199 (55%) | 350 | 286 (81%) |
| *Ci. arietinum* | *Vi. vinifera* | 339 | 201 (59%) | 521 | 414 (79%) |
| *Me. truncatula* | *Vi. vinifera* | 430 | 184 (43%) | 648 | 520 (80%) |
| *Gl. uralensis* | *Vi. vinifera* | 403 | 170 (42%) | 482 | 376 (78%) |
| *Lo. japonicus* | *Vi. vinifera* | 428 | 177 (41%) | 195 | 154 (78%) |
| *Lu. angustifolius* | *Vi. vinifera* | 223 | 73 (32%) | 1654 | 1114 (67%) |
| *Ar. ipaensis* | *Vi. vinifera* | 365 | 171 (47%) | 336 | 275 (81%) |
| *Ar. duranensis* | *Vi. vinifera* | 363 | 152 (42%) | 410 | 327 (79%) |
| *Ch. fasciculata* | *Vi. vinifera* | 426 | 144 (34%) | 186 | 95 (51%) |
| *Ar. thaliana* | *Vi. vinifera* | 392 | 226 (58%) | 707 | 512 (72%) |
| *Vi. vinifera* | *Aq. coerulea* | 478 | 295 (62%) | 303 | 285 (94%) |
| *Aq. coerulea* | *Am. trichopoda* | 552 | 177 (32%) | 130 | 65 (50%) |
| *Am. trichopoda* | *Se. moellendorffii* | 560 | 0 (0%) | 14 | 0 (0%) |

Abbreviations: Segmental duplicates (SD).

The significant number of syntenic SD TFs derived mostly by gene retention after successive WGD events (Table 3). Gene duplicability, the ability of a duplicate pair to remain duplicate, is non-random and biased towards specific gene families, including TFs (Lynch and Conery, 2000;Davis and Petrov, 2004;Li et al., 2016). We analyzed TF duplicability using *Gl. max* TFs from syntenic blocks that survived the 58 mya and the 13 mya WGD events. We used intra-species collinear blocks to identify *Gl. max* SD TFs that correspond to single syntenic regions in *Ph. vulgaris*. We used a maximum Ks threshold of 0.4 to filter the *Gl. max* SD pairs that likely emerged in the 13 mya WGD (Schmutz et al., 2010). Nearly 81% (1808/2230) of the *Gl. max* SD TFs within that Ks range had a syntenic gene in *Ph. vulgaris*. Further, 75% (676/904) of such *Ph. vulgaris* orthologs had a single *Vi. vinifera* syntenic ortholog. In both cases we found that bHLH family had the highest number of syntenic gene pairs (*Gl. max*-*Ph. vulgaris:* 99 pairs and *Ph. vulgaris-Vi. Vinifera:* 43 pairs). Conversely, only 16% (15/93) of the *Gl. max* syntenic singleton TFs correspond to single genomic regions in *Ph. vulgaris* and *Vi vinifera*. We hypothesize that these genes not only depend on the conservation of a local genomic context, but are also sensitive to gene dosage. Our results clearly illustrate the high duplicability of most TF families in soybean and further support the impact of two WGD events that account for a prominent fraction of the TF repertoire of this species.

### Large scale duplication events correlate with increase in TF copy number

Many TF families explored here are broad and diversified, often comprising multiple sub-groups, such as bHLH (Pires and Dolan, 2010), MYB (Du et al., 2012), and ERF (Nakano, 2006). To obtain an overview of the diversification of plant TF families, we assigned them to OGs by using all-vs-all reciprocal BLASTP search, followed by Markov clustering (see methods for details). For example, AP2 had 28 clusters (Supplementary Table S5), labeled as AP:1 to AP:28. We found 1557 TF OGs from the 58 TF families reported above. Nearly 9% (144/1557) of these OGs had no members from legume species, whereas 29% (452/1557) were legume-specific, and 43% (672/1557) had genes from at least 10 species. Expectedly, larger families had more OGs, such as bHLH, C2H2, and MYB, with more than 100 OGs each. Conversely, a few families diverted from this trend, such as SAP (65 genes and 7 OGs) and EIL (143 members and 13 OGs) (Supplementary Figure S4).

To investigate the evolution of TF families in more detail, we analyzed the number of genes per OG in each species using CAFE (v.4.2) (Bie et al., 2006;Han et al., 2013), an elegant method that uses gene birth (λ) and death (μ) rates to model gain and loss events in different lineages of a given ultrametric species tree (see methods for details). We searched for optimal λ and μ based on the maximum likelihood score, using the option to take potentially fragmented genomes into account (see methods for details). We used 672 TF OGs with sufficient variation in number of genes per species (statistical variance ≥ 0.5) and containing genes from at least 10 species. For example, 10 out of 31 AP2 clusters were used for rate estimation (Supplementary Table S5). We repeated the rate parameter search for 50 times and the parameters resulting in the best maximum likelihood score were used for further analysis. The results obtained with CAFE largely confirm the general trend for TF gain upon WGD (Figure 4), which is in line with the above-mentioned correlation between SD, the retention of paralogous TF pairs, and intraspecies synteny. This trend can be exemplified by the nodes representing the legume and *Glycine* ancestors, which have a high number of expanded TF families (Figure 4).

**Figure 4:**
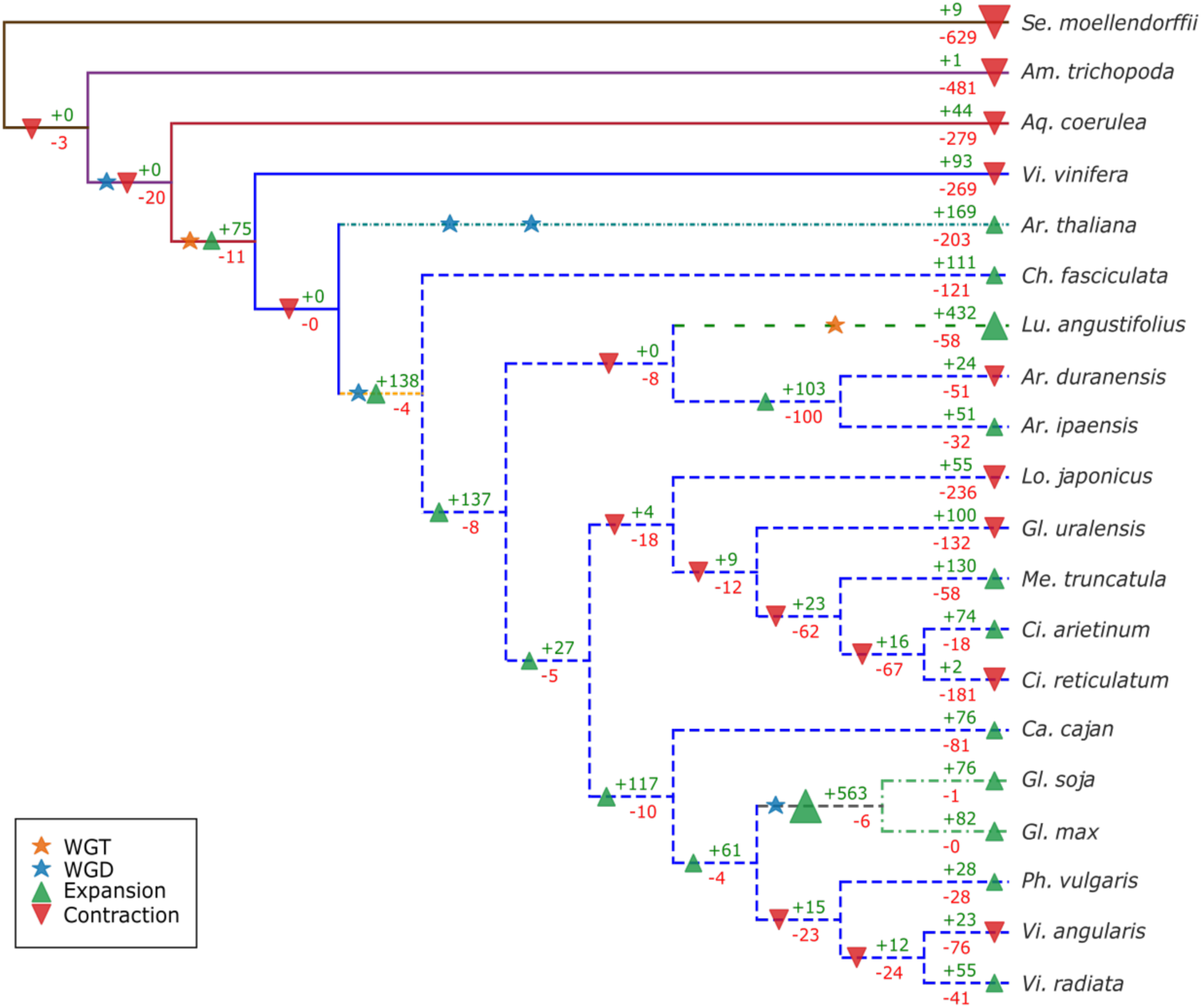
Species tree showing number of transcription factor orthologous groups that gained or lost genes. We used different rates of evolution in different lineages, which are represented as branch styles/colors. Known polyploidization events are marked with stars. Green and red triangles refer to nodes with more expansions and contractions, respectively. Numbers of expanded and contracted orthologous groups are shown in green and red, respectively.

We also investigated the impact of the legume and *Glycine* WGDs in the TF repertoires of *Gl. max* and *Gl. soja*. Firstly, we analyzed the 138 OGs (from 34 TF families) that expanded in legumes in comparison to non-legumes (Figure 4). If all 1557 OGs are considered, an average of 0.21 genes were gained per OG in legumes, in contrast to 1.09 genes in the 138 expanded OGs. Secondly, we identified rapidly evolving OGs (10 of 138; 7.25%) (Table 5), which are those with significant gene gain or loss rate (p-value < 0.05) (Table 6). In these 10 OGs, the average rate of gene gain was 2.0, nearly two and 10 times of that observed in the 138 expanded OGs and in the complete set of 1557 OGs, respectively.

**Table 5:** Number of genes and prevalence of modes of duplication in TF orthologous groups that expanded in legumes.

| Species | Total | SD | TD | PD | rTE | dTE | DD |
| --- | --- | --- | --- | --- | --- | --- | --- |
| *Ca. cajan* | 38 | 14 | 3 | 0 | 0 | 7 | 14 |
| *Ph. vulgaris* | 36 | 14 | 6 | 1 | 0 | 7 | 8 |
| *Vi. radiata* | 36 | 15 | 4 | 1 | 0 | 12 | 4 |
| *Vi. angularis* | 38 | 14 | 9 | 0 | 0 | 9 | 6 |
| *Gl. max* | 67 | 48 | 5 | 3 | 0 | 10 | 1 |
| *Gl. soja* | 69 | 50 | 2 | 3 | 0 | 13 | 1 |
| *Ci. reticulatum* | 22 | 6 | 0 | 0 | 0 | 7 | 9 |
| *Ci. arietinum* | 24 | 11 | 0 | 2 | 0 | 1 | 10 |
| *Me. truncatula* | 39 | 11 | 4 | 2 | 0 | 12 | 10 |
| *Gl. uralensis* | 31 | 12 | 0 | 0 | 0 | 4 | 15 |
| *Lo. japonicus* | 28 | 6 | 2 | 0 | 0 | 13 | 7 |
| *Lu. angustifolius* | 60 | 42 | 6 | 0 | 0 | 0 | 12 |
| *Ar. ipaensis* | 31 | 11 | 4 | 1 | 0 | 9 | 6 |
| *Ar. duranensis* | 32 | 9 | 5 | 1 | 0 | 10 | 7 |
| *Ch. fasciculata* | 34 | 2 | 6 | 0 | 0 | 13 | 13 |

Abbreviations: Segmental duplicates (SD); Tandem duplicates (TD); Proximal duplicates (PD); Retrotransposon mediated duplicates (rTE); Transposon mediated duplicates (dTE); Dispersed duplicates (DD).

**Table 6:**
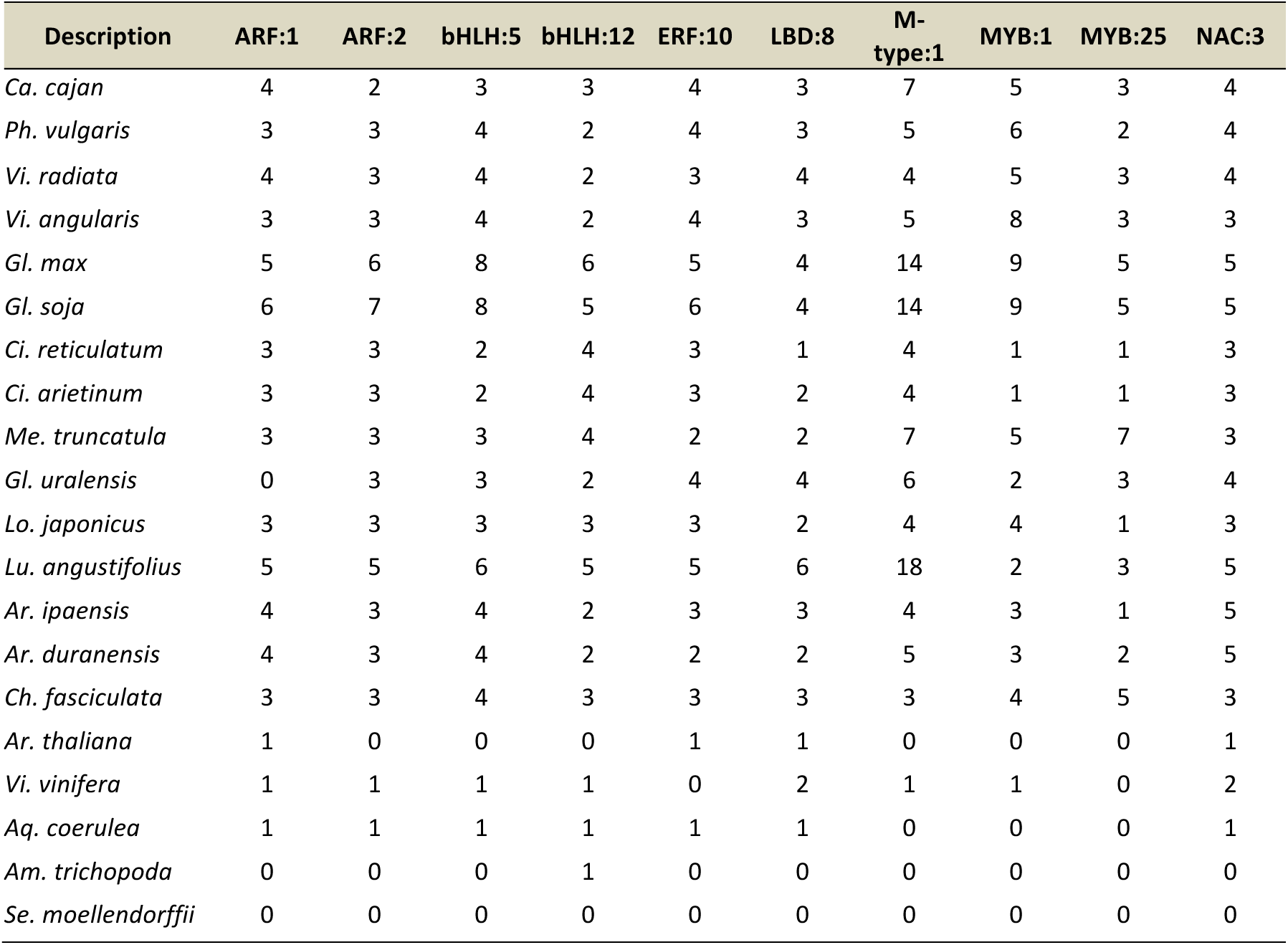
Orthologous groups with significantly (p-value < 0.05) rapid expansion in legumes.

We analyzed the most prevalent modes of duplication in the 138 OGs that expanded in legumes and found that a major fraction of them emerged via SD. While SDs comprise more than 80% of the TFs in species with more recent WGDs (i.e. *Gl. max, Gl. soja*, and *Lu. angustifolius*) (Figure 5), several SD pairs might have lost collinearity after the legume WGD and were classified as DDs. When inspecting the Ks distributions of the paralogous pairs from the 138 OGs that expanded in legumes, we found ranges corresponding to both, the legume and Glycine WGDs (Figure 5) (Schmutz et al., 2010;Cannon, 2013), suggesting that a fraction of the TFs that expanded in the legume WGD subsequently duplicated for a second time in the Glycine WGD. Deviations from this range were observed for *Me. truncatula, Arachis* spp., *Cicer* spp. and, *Ch. fasciculata*, as previously reported (Cannon et al., 2010;Varshney et al., 2013;Tang et al., 2014;Chen et al., 2016). DD paralogs have a more dispersed Ks distribution than that SD, although their Ks distributions also indicate that several DD pairs were likely generated by SD with subsequent loss of collinearity (Figure 5). Collectively, these results support the association between the expansion of legume-specific TF expansions and the WGD event that took place 58 mya.

**Figure 5:**
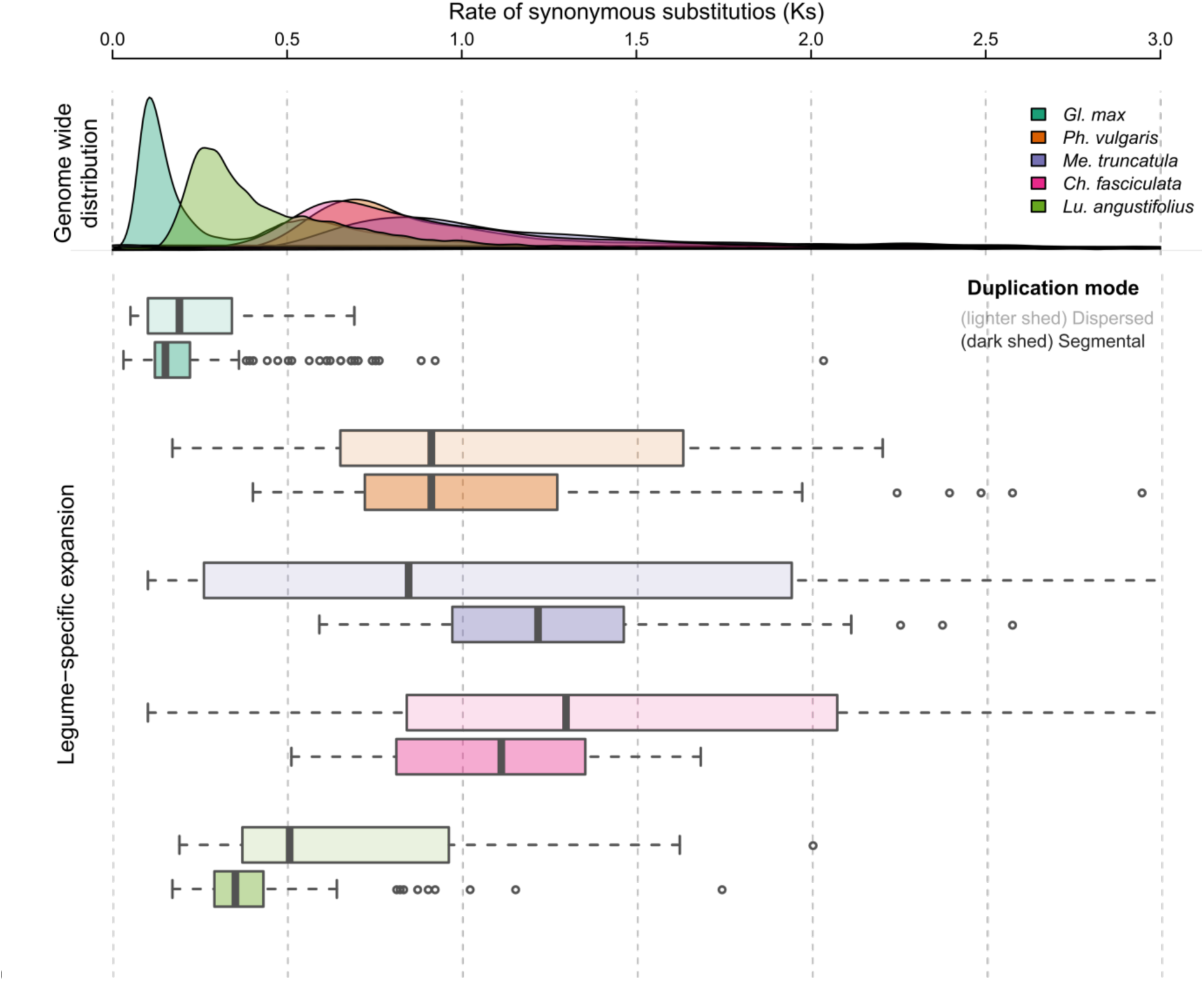
Distribution of synonymous substitution rates (Ks) of 138 orthologous groups with gene gain in legumes. Genome-wide Ks distributions are shown as density plots on the top panel. The bottom panel shows Ks distributions of segmental and dispersed duplicate gene pairs.

To further explore the functional relevance of legume TF expansions, we analyzed gene expression patterns (see methods for details) across multiple tissues from *Me. truncatula*, *Ph. vulgaris*, and *Gl. max* (Figure 6; Supplementary Figure S5). Strikingly, the 138 OGs that expanded in legumes are enriched in genes with preferential expression in nodules (Fisher’s Exact Test, p-values = 1.7 × 10^-4^ and 1.4 × 10^-5^ for *Me. truncatula* and *Gl. max*, respectively) and roots (Fisher’s Exact Test, p-values = 1.4 × 10^-9^ and 5.9 × 10^-3^ for *Me. truncatula* and *Ph. vulgaris*, respectively). These results indicate that the recruitment of these genes predate the emergence of nodulation in legumes and might have played roles in the root physiology associated with this process.

Next, we integrated phylogenetic reconstructions of the 10 rapidly expanded OGs with gene expression data and found three interesting groups (i.e. bHLH:12, M-type:1, ERF:10) (Table 6). The bHLH:12 OG showed significantly higher expression during nodule development in *Me. truncatula, Ph. vulgaris*, and *Gl. max* (Figure 6B and Supplementary Figure S5). This OG includes two SD pairs of *Me. truncatula* bHLHs, Medtr4g087920-Medtr2g015890 and Medtr4g079760-Medtr2g091190 with Ks values of 1.0523 and 0.8292, respectively. *Ph. vulgaris* and *Gl. max* orthologs of these genes were also more expressed in roots and nodules than in other tissues (Figure 6).

**Figure 6:**
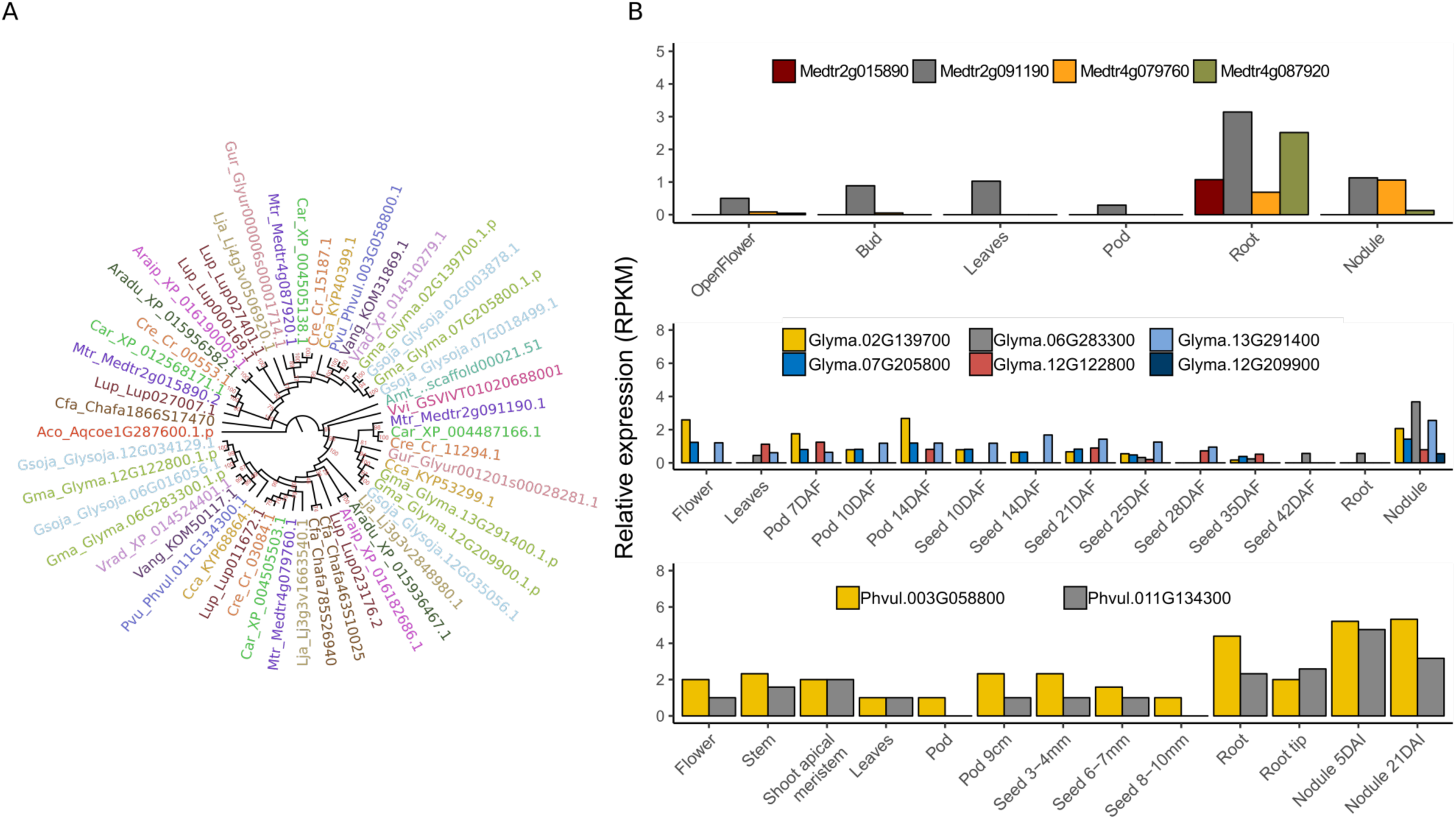
A. Phylogenetic reconstruction of the orthologous group bHLH:12, which is expanded in legumes. B. Gene expression patterns of bHLH:12 genes in *Me. truncatula* (BioProject: PRJNA80163)*, Gl. max* (Libault et al., 2010) and *Ph. vulgaris* (O’Rourke et al., 2014), showing a trend for greater expression in roots and nodules.

Another interesting OG encodes TFs from the M-type MADS family (M-Type:1). This OG has independently expanded in different species (Supplementary Figure S6), including a major expansion in *Lu. angustifolius*. Nine of 14 *Gl. max* genes in this cluster emerged within the Glycine genus, including one gene (Glyma.03G083700.1) with preferential expression in seeds and flowers (Supplementary Figure S7A). The two *Ph. vulgaris* orthologs (Phvul.006G077700 and Phvul.006G077800.1) showed seed-specific expression, suggesting their importance in seed development (Supplementary Figure S7B). Interestingly, these *Ph. vulgaris* genes originated from an ancestral tandem duplication, as this organization is also found in other legumes (Supplementary Figure S6). Although this OG lacked an *Ar. thaliana* member, the closest *Ar. thaliana* homologs include *AT5G27810*, *AT1G22590* (AGAMOUS-LIKE 87, *AGL87*), and *AT5G48670* (AGAMOUS-LIKE80, *AGL80*). Importantly, *AGL80* has been shown to be responsible for central cell and endosperm development in *Arabidopsis* (Portereiko et al., 2006).

We also analyzed two ERF (*ethylene response factor*) OGs (ERF:10 and ERF:18) containing genes playing critical roles in nodulation. Of these two, only ERF:10 was among the 10 rapidly expanded OGs. Manual curation revealed that ERF:10 and ERF:18 comprise ERF *required for nodule differentiation* (EFD) and ERF *required for nodulation* (ERN) genes, respectively. ERN and EFD genes regulate nodulation in *Me. truncatula* (Vernie et al., 2008;Young et al., 2011;Cerri et al., 2012). Three of the four *MtERNs* (i.e. Medtr7g085810.1, Medtr6g029180.1, and Medtr8g085960.1) had relatively higher expression after inoculation than in roots or nodules, supporting their critical role in nodule development (Supplementary Figure S9A). The biased expression towards nodules and root tissues are also observed in *Ph. vulgaris* and *Gl. max* orthologs (Supplementary Figure S8). Of the two *MtEFDs*, Medtr4g008860 and Medtr3g106290 were more expressed in nodules and inoculated root hairs, respectively. Similarly, *Ph. vulgaris* and *Gl. max* ERNs are also more expressed in roots and nodules than in aerial tissues (Supplementary Figure S8).

### Association between quantitative traits and *Glycine max* specific TFs

The *Glycine* node had the largest number of expanded OGs, with 36% (563/1557) (Figure 4) of them expanding by an average rate of 1.67 genes per OG. Out of these, 57 had rapidly expanded (p-value < 0.05) with an average rate of 2 genes per OG. In comparison to the *Glycine* node, 76 and 82 OGs expanded in *Gl. soja* and *Gl. max*, respectively. Of the OGs that expanded in *Gl. max*, 79% (65/82) showed significant expansions (p-value < 0.05), with average rate of 1.44. Among these families, ERF (6 OGs), MYB (5 OGs), MYB_related (8 OGs), bHLH (7 OGs), and C2H2 (7 OGS) TFs gained more than 5 genes per OG. Interestingly, 59% (202/341) of the genes from expanded OGs lie within SD regions, supporting the importance of the Glycine WGD in shaping these families in *Gl. max*.

We explored whether some of these OGs could be related with important soybean agronomic traits. We searched the *Gl. soja* syntenic regions corresponding to the 341 *Gl. max* TFs from the 65 rapidly expanded OGs. We identified 50 TFs without a homeolog in *Gl. soja* (Supplementary Table S6), out of which only five had a syntenic ortholog in *Ph. vulgaris*. Interestingly, two ERF (Glyma.20G115300, Glyma.14G161900) and one SBP (Glyma.06G205700) TFs are within previously reported chromosomal regions associated with important quantitative traits (Supplementary Figure S9) (Fang et al., 2017). The ERF Glyma.20G115300 was located within a region associated with overall leaf size and average number of seeds per pod. The second ERF, Glyma.14G161900, is within a region associated with FA18 content and ratio in mature seeds. Finally, the SBP TF Glyma.06G205700 is within a region regulating branch density (i.e. ratio of branch number and plant height) and beginning bloom date. (Supplementary Table S6). We envisage that many more of these 50 TFs will be associated with important traits, which could be revealed in by a more comprehensive work integrating QTL information from other genotypes and studies with our phylogenomic results.

## Methods

### Genomic data

Genome sequences and annotations for 20 plant species were obtained from public repositories (Table 1). We used coding sequences and predicted protein sequences from the longest splicing isoform (when more than one are available). Predicted proteins with less than 50 amino acids or containing premature stop codons or more than 20% ambiguous amino acids were excluded.

### Prediction and classifications of transcription factors (TFs)

To remove redundancy due to splicing isoforms and incomplete gene predictions, we removed nearly identical sequences using BLASTCLUST (Altschul et al., 1997) as previously described (parameters: -S 1.89 -L 0.9 -b F) (Gossani et al., 2014;Vidal et al., 2016).

We adopted the TF family classification scheme of plantTFDB (Zhang et al., 2011;Jin et al., 2017). We created a local database of protein domains by combining all HMM profiles from PFAM-A (Release 31.0) (Finn et al., 2016) and 13 plant specific TF HMM profiles downloaded from PlantTFDB (Supplementary Table S1). Protein sequences were searched for conserved domains using HMMER 3.0 (http://hmmer.org) (domain e-value cutoff < 0.01). TFs were classified in 58 families according to their DBD.

### Species phylogeny

A species phylogeny was reconstructed using low copy orthologs present in all 20 species. We clustered the predicted proteins on the basis of the pairwise sequence similarity of their longest protein products, which was computed with BLAST (e-value ≤ 1e^-5^) (Altschul et al., 1997). Sequence pairs with percentage identity of at least 35% and query coverage of at least 50% were used for Markov clustering using mclblastline (v. 12-068; Inflation parameter: 1.5) (Enright et al., 2002). Clusters containing up to 22 genes with at least one gene from each species were used. If a species had paralogous genes, the paralog with greater identity to orthologs from other species was used. Amino acid sequence alignment was performed using DECIPHER (Wright, 2015) and cDNA alignment performed with PAL2NAL (Suyama et al., 2006). We concatenated the codon alignments of these genes to create a super-alignment. Next, phangorn (Schliep, 2011) was used to estimate the best substitution model for the phylogenetic reconstruction, which was performed using RAxML (v8.2.11; model: GTRGAMMAIG4, bootstrap: 1000) (Stamatakis, 2014). The phylogram and sequence alignment were used in relTime-ML (implemented in MEGA-X, v.10.0.1) (Tamura et al., 2018) to generate an ultrametric tree. We used the TimeTree database (Kumar et al., 2017) to retrieve the divergence times of Fabaceae and *Vi. vinifera* (110 mya) and of *Ph. vulgaris* and *Gl. max* (24 mya), which were used as references.

### Synteny and synonymous mutation rate (Ks)

We identified segmental duplications using DAGCHAINER (r02062008) (Haas et al., 2004) by using bidirectional best BLASTP hits (e-value ≤ 1e^-10^, 35% minimum identity, 50% minimum query coverage). A minimum of four collinear genes were required to identify a syntenic block (DAGCHAINER, parameter -A 4), as previously used in soybean (Severin et al., 2011). Tandem duplicates were also identified using DAGCHAINER (parameters -T -A 2). Tandem or segmental gene pairs had their non-synonymous (Ka) and synonymous (Ks) mutation rates estimated using the *bp_pairwise_kaks* script, distributed with BioPerl (v5.22.1) (Stajich et al., 2002).

### Orthologous cluster and TF paralog identification

TF OGs were inspected for different modes of gene duplication. Multiple modes of duplication can also co-occur in a group of genes. In these cases, only one type of duplication is reported, following the order SD>TD>PD>rTE>dTE>DD (Proulx et al., 2011;Qiao et al., 2018). Although this strategy can slightly underestimate some duplication levels, it helped us to assess the main forces shaping the expansion of TF families.

### Estimation of expansions and contractions in TF families

We used CAFE (v4.2) (Han et al., 2013) to assess the evolution of TF family sizes using the time-calibrated species tree and TF OG compositions as inputs. We used the *cafeerror.py* script, available in the CAFE package, to model error rates that might have been introduced in gene family sizes, particularly by species with more fragmented genome assemblies (e.g. *Lu. angustifolius*) (Han et al., 2013). This error model was used adjust family sizes. We estimated λ and μ by running CAFE for 50 times and selected the parameters that gave the best maximum likelihood estimate. These parameters were used to estimate OG sizes at ancestor nodes and to predict rapidly evolving OGs (p-value < 0.05), which are those that significantly gained or lost genes. The average net change at each node on the species tree was expressed as:

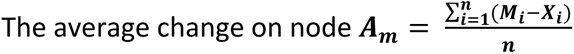

where *n* is the total number of OGs, (*M_i_*−*X_i_*) is the difference in OG size between node *M* and its parent node *X* for a given OG *i*. A negative or positive *A_m_* value stands for contraction or expansion, respectively. Some remarkably expanded or contracted TF OG had their phylogenies reconstructed with RAxML (v8.2.11; model: GTRGAMMAAUTO, bootstrap: 1000) and visualized using Figtree (v.1.4.3) (http://tree.bio.ed.ac.uk/software/figtree/).

### Gene expression data and tissue specificity

*Ar. thaliana* and *Ph. vulgaris* normalized gene expression data were obtained from public databases such as ArrayExpress (Liu et al., 2012) and *Pv*GEA (O’Rourke et al., 2014), respectively. Four additional RNAseq datasets were downloaded from the NCBI SRA database (https://www.ncbi.nlm.nih.gov/sra/). The first two datasets comprise two soybean transcriptome studies (Bioproject PRJNA208048, PRJNA79597) (Libault et al., 2010;Severin et al., 2010). The third dataset includes *Me. truncatula* transcriptomes (Boscari et al., 2013) in the following conditions and tissues: nitrogen-starving roots, roots inoculated with *Sinorhizobium meliloti*, and root nodules (BioProject PRJNA79233). We also downloaded an additional *Me. truncatula* RNAseq data covering 7 different tissues (BioProject PRJNA80163). RNAseq reads were mapped on each species genome using STAR v2.5.3a (Dobin et al., 2013) and normalized gene expression values were estimated with StringTie v1.3.4d (Pertea et al., 2015), both with default parameters.

Expression values lower than 1 were converted to 0 and considered not expressed. We added 1 to all values, which were then log_2_ transformed. To determine tissue-preferential expression, we transformed the gene expression values in a transformed z-score index (Kryuchkova-Mostacci and Robinson-Rechavi, 2016). Depending on the highest expression in a given tissue, genes with transformed z-score index > 0.9 were labeled as preferentially expressed.

### Micro-syntenic regions between *Gl. max*, *Gl. soja*, and *Ph. vulgaris*

We used DAGCHAINER output files to identify the microsyntenic regions in *Gl. max, Gl. soja*, and *Ph. vulgaris*. In particular, we queried the genes from OGs with significantly larger (as predicted by CAFE) sizes in *Gl. max* in comparison to the *Glycine* node. For each *Gl. max* gene, we considered only one collinear region from *Gl. soja* and *Ph. vulgaris*. When more than one collinear region was detected, we selected that with the highest DAGCHAINER alignment score. We visualized the microsynteny regions using Genome Context Viewer available on Legume Information System (Cleary et al., 2017).

### QTL intervals

We obtained the chromosomal coordinates of 150 QTLs significantly associated with 57 soybean traits (Fang et al., 2017). Chromosomal coordinates of soybean genes were mapped to these QTL regions using bedtools v2.26.0 (Quinlan and Hall, 2010).

## Acknowledgements

This work was supported by funding from Coordenação de Aperfeiçoamento de Pessoal de Nível Superior (CAPES, Finance code 001), Conselho Nacional de Desenvolvimento Científico e Tecnológico (CNPq), and Fundação Carlos Chagas Filho de Amparo à Pesquisa do Estado do Rio de Janeiro (FAPERJ).

