## Supplementary figures and tables S1, S3 and S5 for "Polyploidization events shaped the transcription factor repertoires in legumes (Fabaceae)"

Laboratório de Química e Função de Proteínas e Peptídeos, Centro de Biociências e Biotecnologia, Universidade Estadual do Norte Fluminense Darcy Ribeiro; Campos dos Goytacazes, Brazil.

* Corresponding author

Av. Alberto Lamego 2000 / P5 / 217; Parque Califórnia

Campos dos Goytacazes, RJ

Brazil

CEP: 28013-602

### Supplementary figures

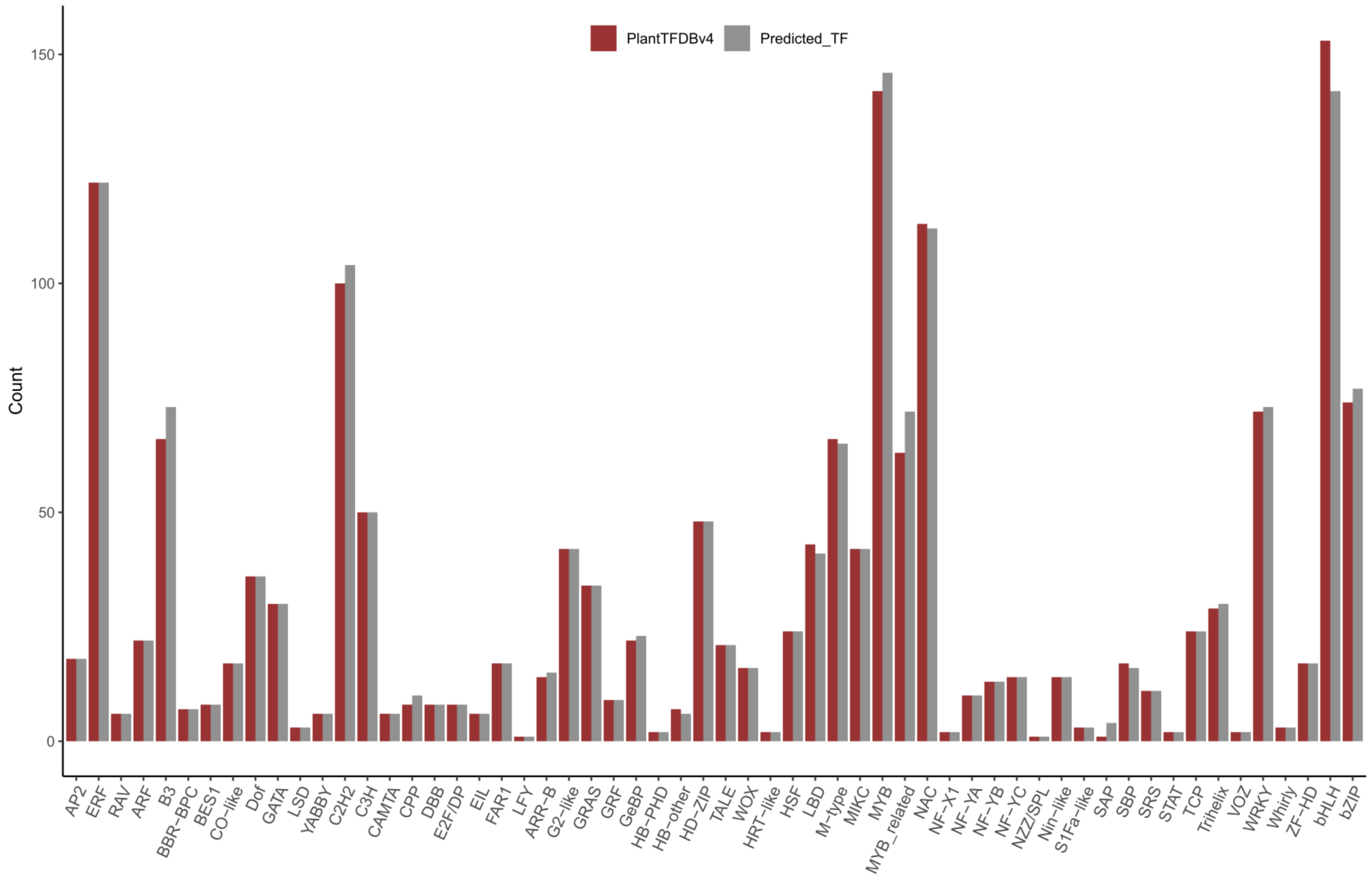


Supplementary Figure S1: Number of *Arabidopsis thaliana* transcription factors predicted using our pipeline in comparison with that available in PlantTFDB.

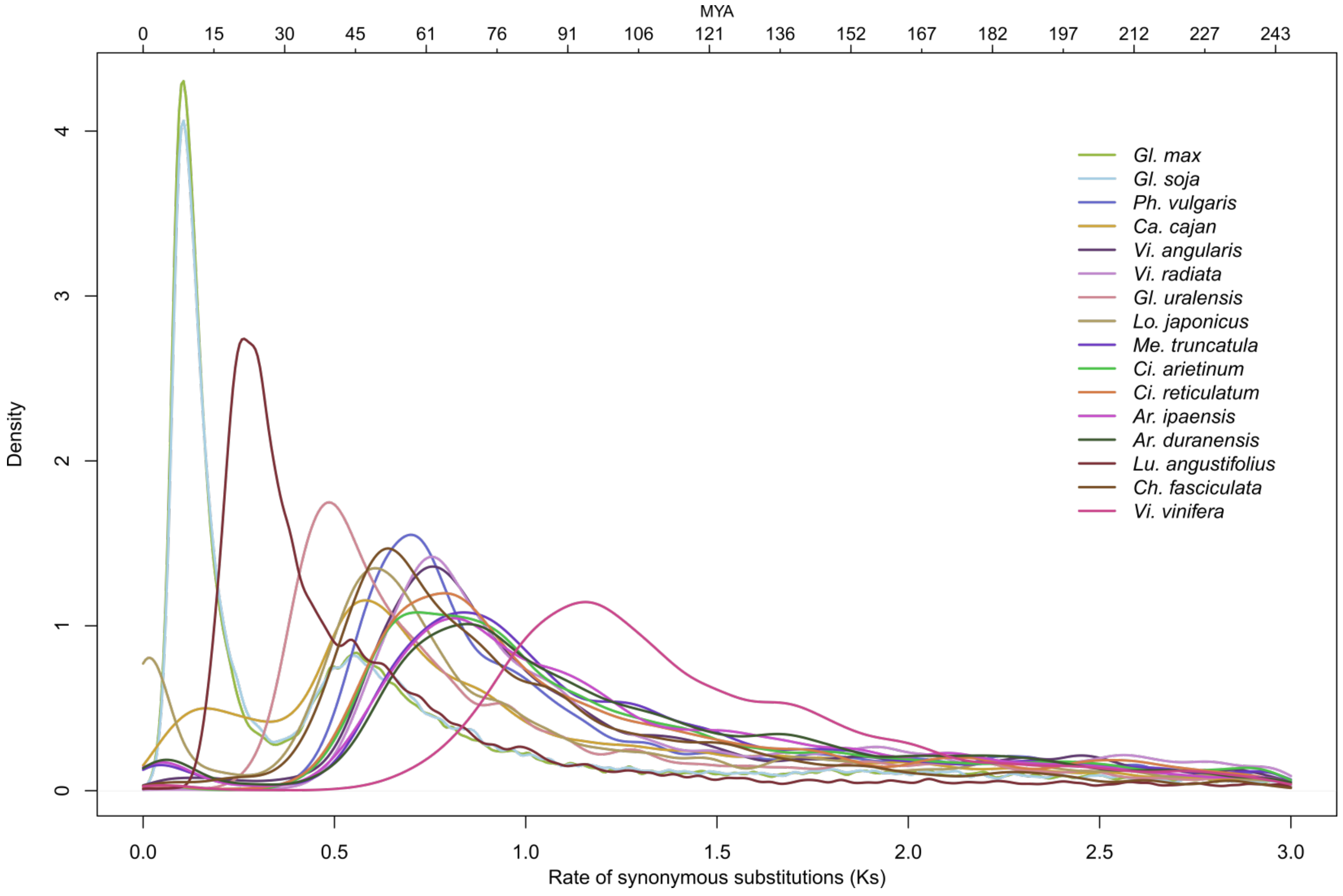


Supplementary Figure S2: Ks distribution of collinear paralogous pairs in *Vitis vinifera* and legumes. Ks peaks from each species likely represent whole-genome duplications. The time scale at the top was used to estimate the age of the duplication event and was estimated as T=Ks/2λ, where λ is number of synonymous substitutions per synonymous substitutions sites per year. We estimated T using λ =6.1*1e-9, as previously described (Lynch and Conery, 2003).

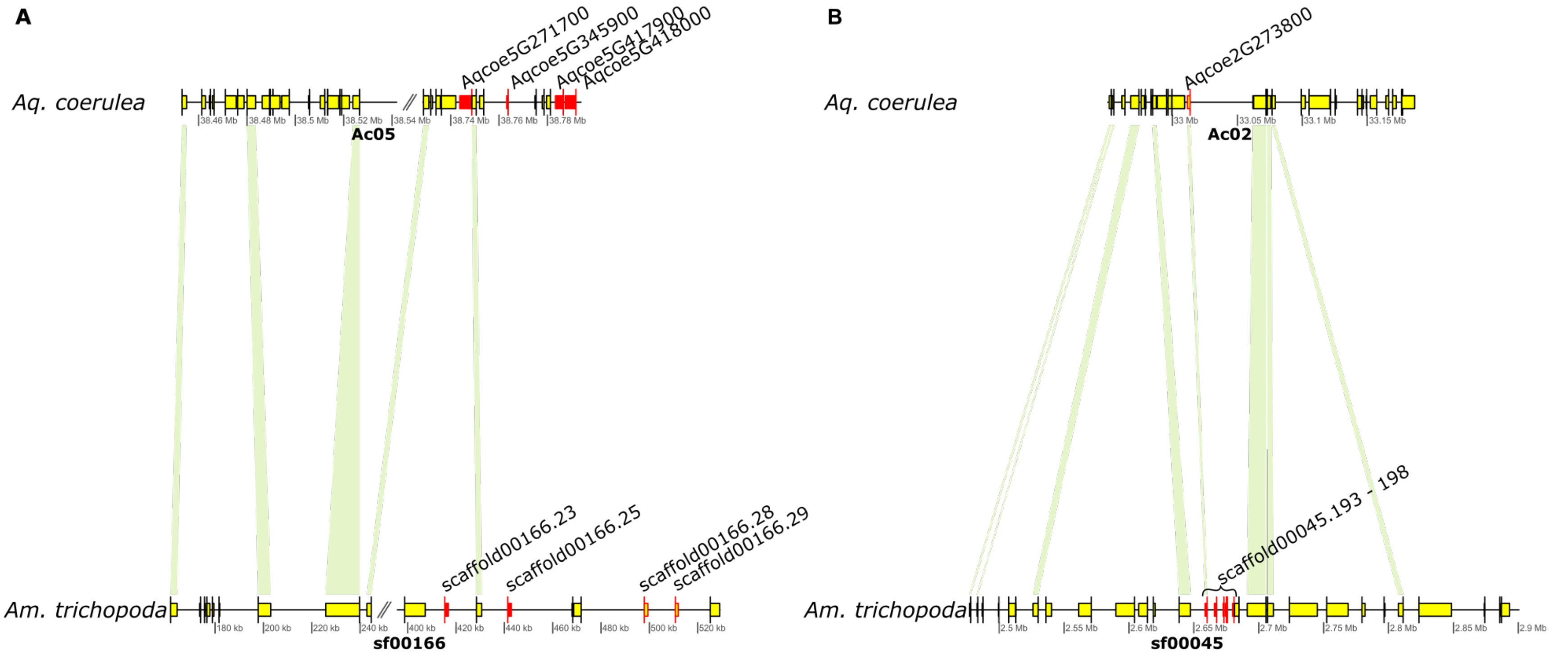


Supplementary Figure S3: Schematic representation of collinear regions in *Aquilegia coerulea* and *Amborella trichopoda*, showing the expansion of GRAS TFs in the latter. In both panels (A and B), the upper and lower bars represent pseudo-chromosomes/contigs from *Aq. coerulea* and *Am. trichopoda*, respectively. Genes are represented by yellow arrows. Green shades connect homologous genes in the two species. Locally duplicated genes are shown as red arrows. GRAS genes are labeled with gene names and have red borders. A. *Aq. coerulea* (Ac05:38.45Mb-38.73Mb) versus *Am. trichopoda* (Sf00166:0.16Mb-0.52Mb), B. *Aq. coerulea* (Ac02:32.95Mb-33.18Mb) versus. *Am. trichopoda* (Sf0045:2.47Mb-2.90Mb).

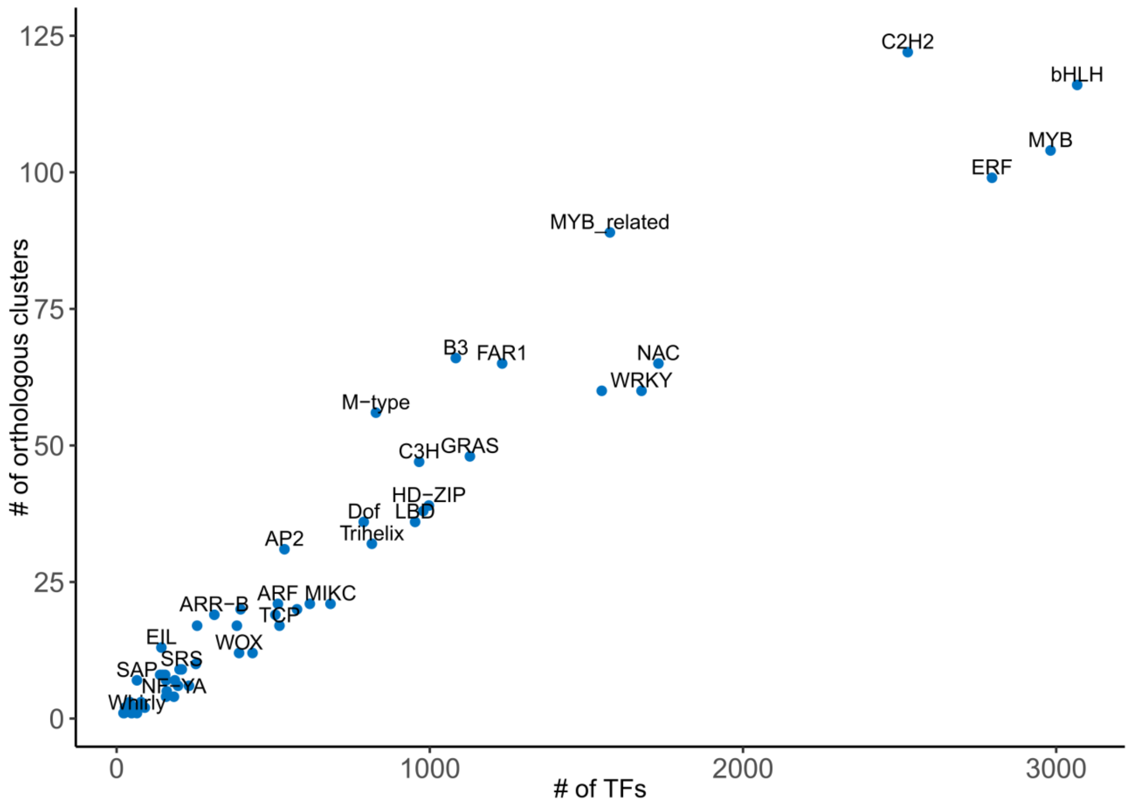


Supplementary Figure S4: Number of orthologous groups and number of members in each transcription factor family.

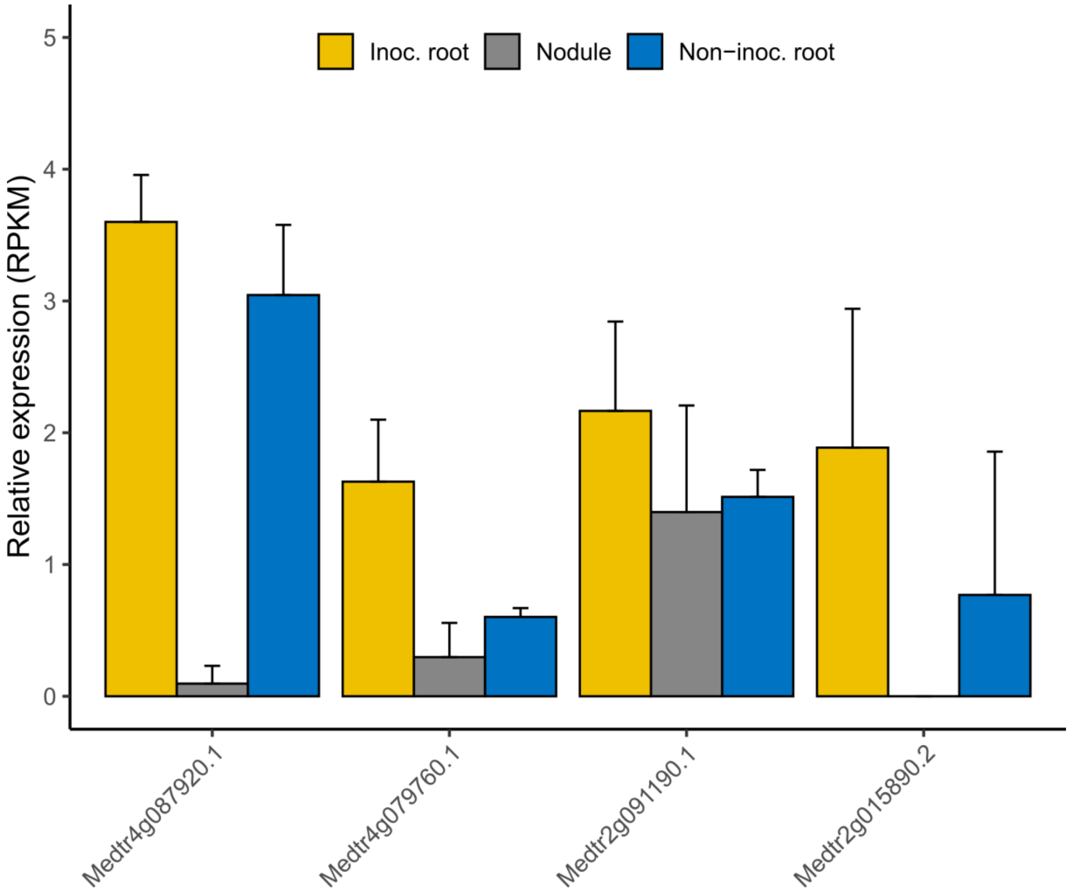


Supplementary Figure S5: Normalized expression of genes from the bHLH:12 orthologous group. Gene expression patterns of four *Me. truncatula* bHLH genes. RNA-seq data were obtained from a previous study (Boscari et al., 2013). Non-inoc. Root: nitrogen-starving roots; Inoc.-root: roots inoculated with *Sinorhizobium meliloti*; Nodule: root nodules.

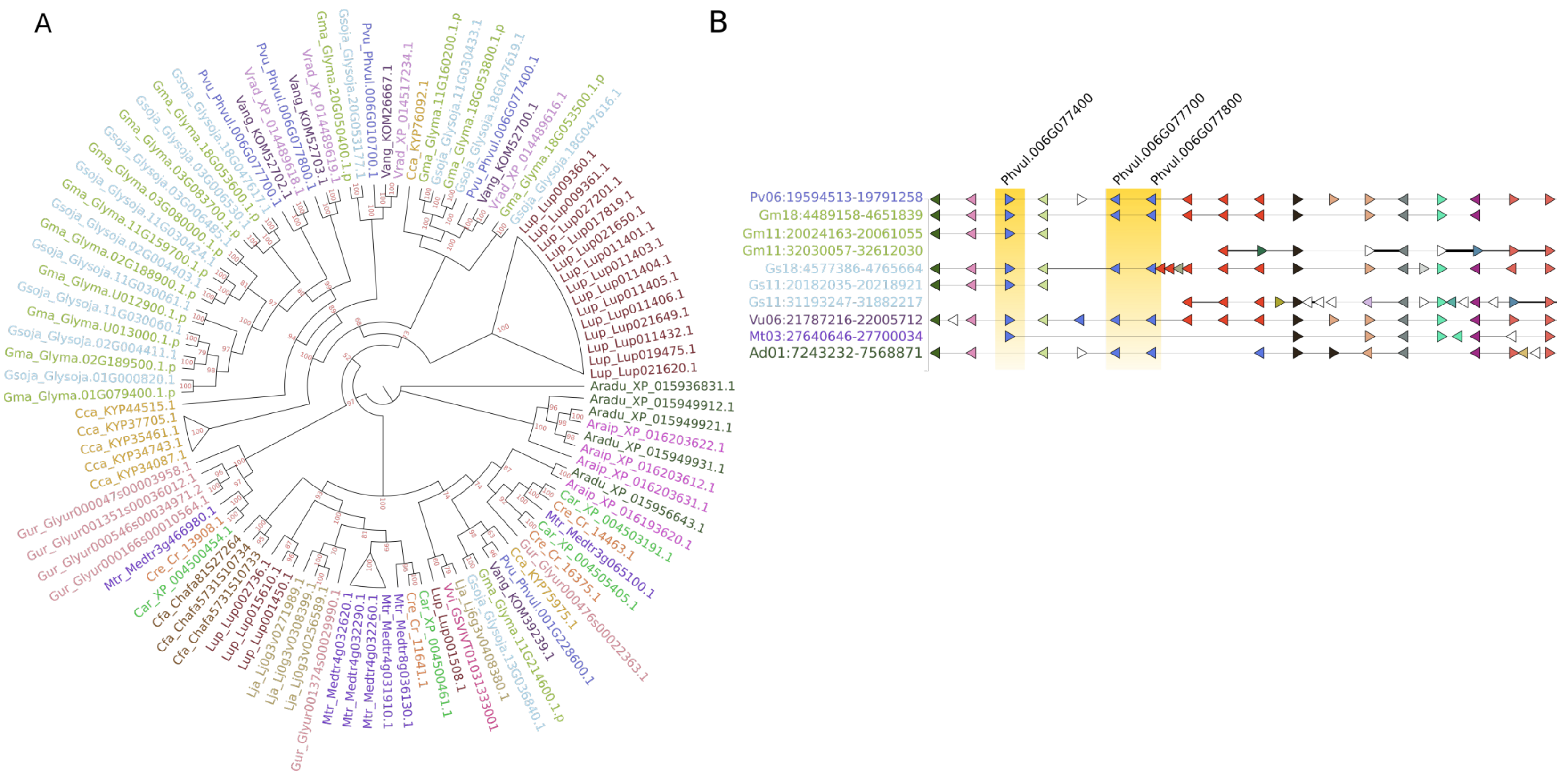


Supplementary Figure S6: Phylogenetic tree of the M-type:1 orthologous group. Genes from same species are labeled with similar colors. Clades comprising genes from the same species were collapsed. The Vigna species used in our analyses are not available in Genome Context Viewer at Legume Information System (Cleary et al., 2017). Hence, we used a close species, *Vigna unguiculata*, to illustrate the syntenic relationships.

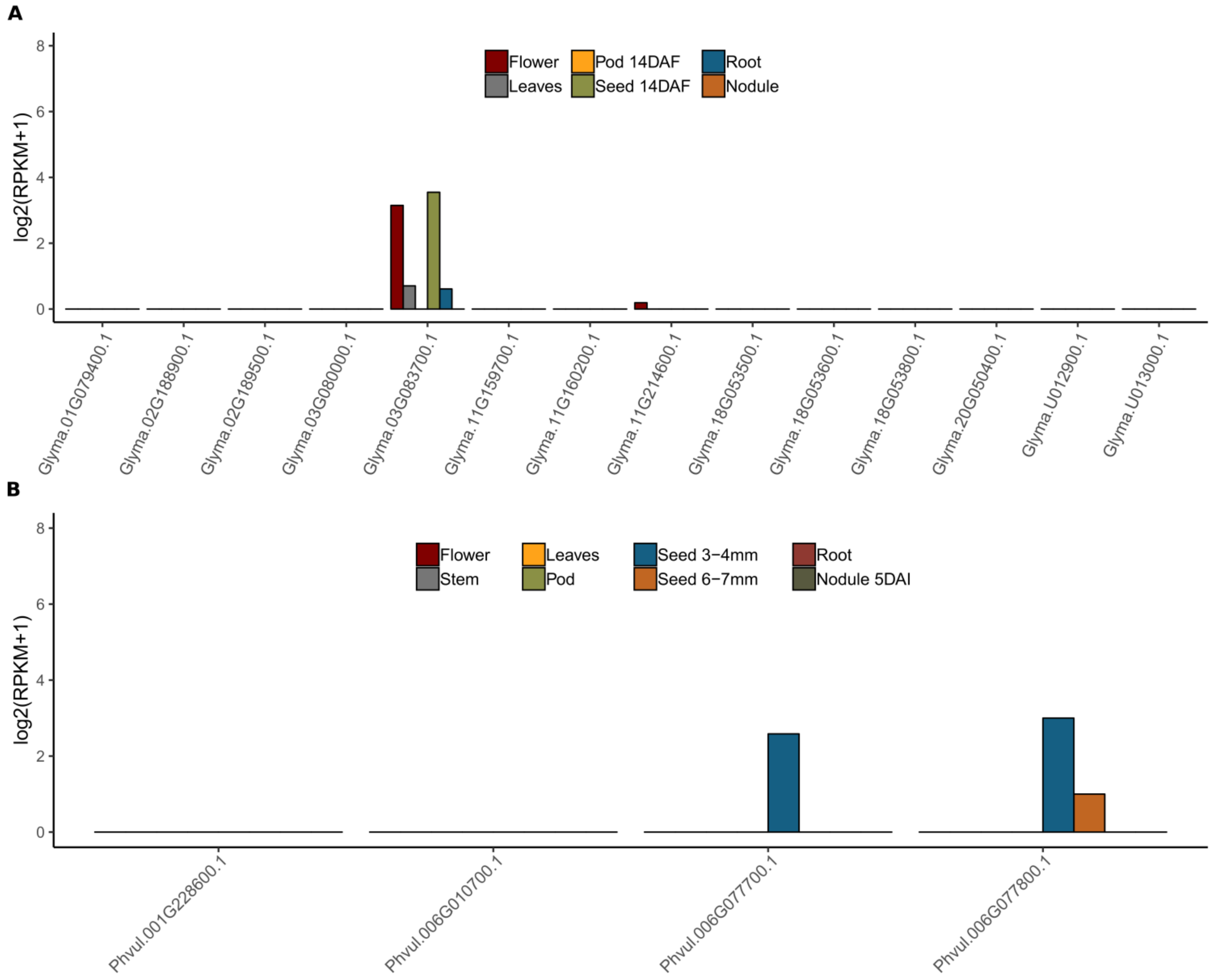


Supplementary Figure S7: Expression levels of M-type:1 genes. A and B represent expression levels of M-type:1 genes across various tissues in *Gl. max* (Severin et al., 2010) (A) and *Ph. vulgaris* (O'Rourke et al., 2014) (B).

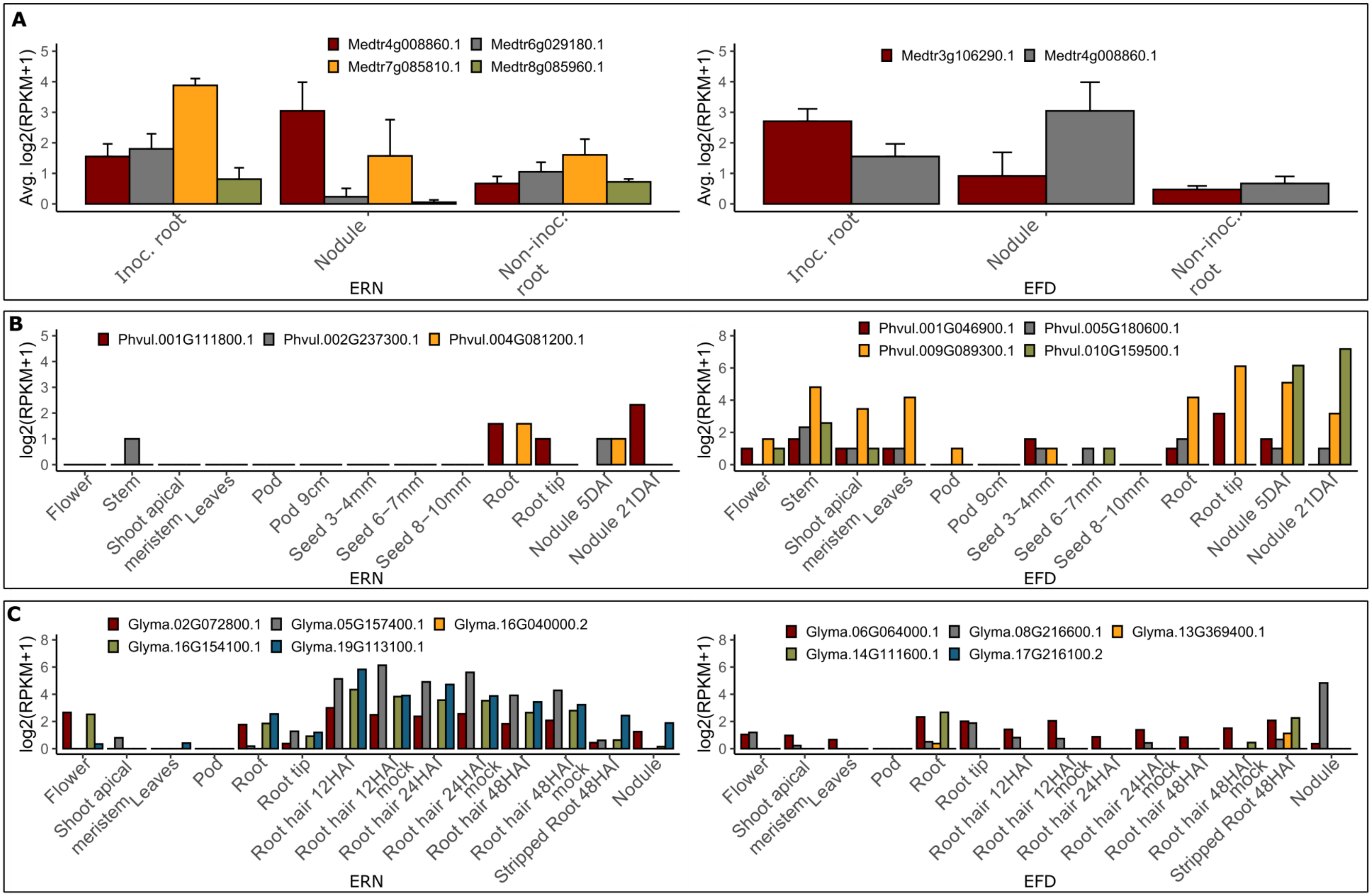


Supplementary Figure S8: Expression levels of ERF required for nodulation (ERN) and nodule differentiation (EFD) genes in *Me. truncatula* (A), *Ph. vulgaris* (B) and *Gl. max* (C). In all three panels, left and right images represent ERN and EFD genes, respectively.

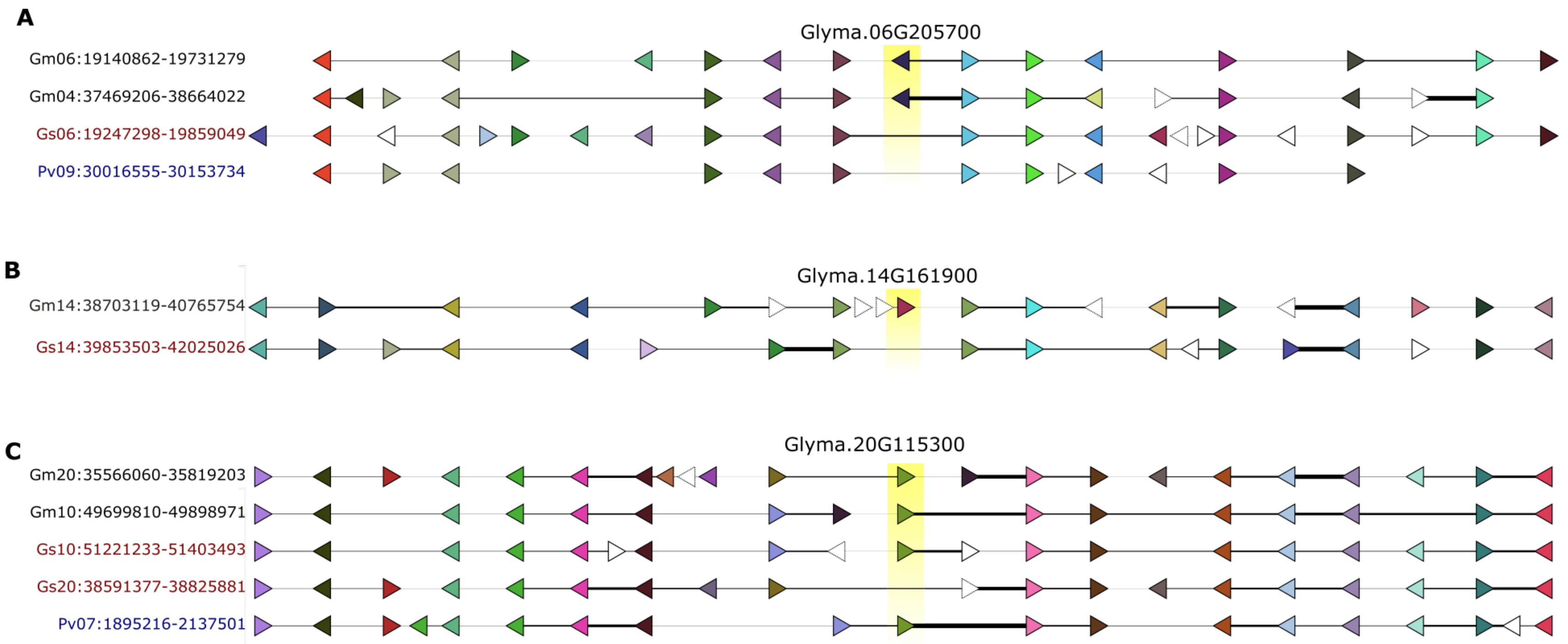


Supplementary Figure S9: Microsynteny of *Gl. max* genes located within chromosomal regions associated with phenotypic traits. The genomic context was rendered on Genome Context Viewer at Legume Information System (Cleary et al., 2017). The queried *Gl. max* genes are on the top rows, labeled and highlighted in yellow. Homologous genes are drawn with the same colors. Singletons are in white. The thickness of the horizontal lines is proportional to the intergenic distances. A. *Glyma.06G205700* is located in a region associated with branch density (i.e. ratio of branch number and plant height); B. *Glyma.14G161900* is within a region associated with fatty acid (FA18) content in mature seeds. C. *Glyma.20G115300* is within a region associated with leaf length, shape, and width, number of four seeds per pod, ratio of four seeds per pod, and ratio of two seeds per pod.

Supplementary Table S1: Conserved domains and rules used for identifying transcription factors. A transcription factor was expected to have at least one DNA binding domain and may contain auxiliary domains. Forbidden domains were used to eliminate false positives.

| Superfamily | Family | Description | DBD* domain [# of domains] (Pfam accession) | Auxiliary domain | Forbidden domain |
|---|---|---|---|---|---|
| AP2/ERF | AP2 | APETALA 2 | AP2[>=2] (PlantTFDB) | | |
| | ERF | Ethylene Responsive Factor | AP2[1] (PlantTFDB) | | |
| | RAV | Related to ABI3/VP1 | AP2[1] (PlantTFDB) and B3(PF02362) | | |
| B3-superfamily | B3 | B3 | B3(PF02362) | | |
| | ARF | Auxin response factor | B3(PF02362) | Auxin_resp (PF06507) | |
| BBR-BPC | BBR-BPC | Barley B Recombinant (BBR) - BASIC PENTACYSTEINE1 (BPC1) | GAGA_bind (PF06217) | | |
| BES1 | BES1 | BRI1-EMS-SUPPRESSOR | BES1_N (PF05687) | | Glyco_hydro_14 (PF01373) |
| bHLH | bHLH | basic helix loop helix | HLH (PF00010) | | |
| bZIP | bZIP | Basic leucine zipper | bZIP_1 (PF00170) | | |
| C2C2 | Dof | DNA binding with one finger | Zf-Dof (PF02701) | | |
| | GATA | GATA | GATA-zf (PF00320) | | |
| | CO-like | CONSTANS-like | Zf-B_box (PF00643) | CCT (PF06203) | |
| | YABBY | YABBY | YABBY (PF04690) | | |
| | LSD | LESION SIMULATING DISEASE 1, LSD1 | Zf-LSD1 (PF06943) | | Peptidase_C14 (PF00656) |
| | C2H2 | CCHH (Zn) | Zf-C2H2 (PF00096) | | Exonuc_X-T(PF00929) |
| C3H | C3H | CCCH (Zn) | Zf-CCCH (PF00642) | | RRM_1 (PF00076) or Helicase_C (PF00271) |
| CAMTA | CAMTA | Calmodulin binding transcription factors | CG1 (PF03859) | | |
| CPP | CPP | Cystein-rich polycomb-like protein | CXC (PF03638) | | |
| DBB | DBB | Double B-box zinc finger | zf-B_box[>=2] (PF00643) | | |
| E2F/DP | E2F/DP | E2 factor protein and DP protein | E2F_TDP (PF02319) | | |
| EIL | EIL | Ethylene-Insentive3 (EIN3)-like protein3 (EIL3) | EIN3 (PF04873) | | |
| FAR1 | FAR1 | FAR-RED IMPAIRED RESPONSE1 | FAR1 (PF03101) | | |
| FLO | LFY | LEAFY | FLO_LFY (PF01698) | | |
| GARP | G2-like | Golden2 (G2)-like | G2-like (PlantTFDB) | Response_reg (PF00072) | |
| | ARR-B | Type-B phospho-accepting response regulator (ARR) family | G2-like (PlantTFDB) | | |
| GeBP | GeBP | GLABROUS1 enhancer-binding protein (GeBP) | DUF573 (PF04504) | | |
| GRAS | GRAS | GAI, RGA, and SCR | GRAS (PF03514) | | |
| GRF | GRF | GROWTH-REGULATING FACTOR (GRF) | WRC (PF08879) | QLQ (PF08880) | |
| HB | HD-ZIP | HD-Zip | Homeobox (PF00046) | HD-ZIP_I/II (PlantTFDB) or SMART (PF01852) | |

| | | | | |
|---|---|---|---|---|
| | TALE | Three Amino acid Loop Extension | Homeobox (PF00046) | BELL (PlantTFDB) or ELK (PF03789) |
| | WOX | WUS homeobox-containing protein family | Homeobox (PF00046) | Wus type homeobox (PlantTFDB) |
| | HB-PHD | HB-PHD finger | Homeobox (PF00046) | PHD (PF00628) |
| | HB-other | HB-other | Homeobox (PF00046) | |
| HRT-like | HRT-like | Hairy-Related transcription-factor-like | HRT-like (PlantTFDB) | |
| HSF | HSF | Heat shock factor | HSF_dna_bind (PF00447) | |
| LBD (AS2/LOB) | LBD | ASYMMETRIC LEAVES2/LATERAL ORGAN BOUNDARIES | LOB (PF03195) | |
| MADS | M-type | MADS-type I | SRF-TF (PF00319) | |
| | MIKC | MADS-type II | SRF-TF (PF00319) | K-box (PF01486) |
| MYB superfamily | MYB | Myb proto-oncogene protein | Myb_dna_bind [1] (PF00249) | SWIRM (PF04433) |
| | MYB_related | Myb-related | Myb_dna_bind [>=2] (PF00249) | SWIRM (PF04433) |
| NAC | NAC | NAM, ATAF1, 2 and CUC2 | NAM (PF02365) | |
| NF-X1 | NF-X1 | Nuclear factor, X-box binding 1 | Zf-NF-X1 (PF01422) | |
| NF-Y | NF-YA | Nuclear factor Y subunit A | CBFB_NFYA (PF02045) | |
| | NF-YB | Nuclear factor Y subunit B | NF-YB (PlantTFDB) | |
| | NF-YC | Nuclear factor Y subunit C | NF-YC (PlantTFDB) | |
| Nin-like | Nin-like | NODULE INCEPTION | RWP-RK (PF02042) | |
| NZZ/SPL | NZZ/SPL | SPOROCYTELESS/NOZZLE | NOZZLE (PF08744) | |
| S1Fa-like | S1Fa-like | S1Fa-like | S1FA (PF04689) | |
| WRKY | WRKY | WRKY | WRKY (PF03106) | |
| Trihelix | Trihelix | Trihelix | Trihelix (PlantTFDB) | |
| TCP | TCP | TEOSINTE-LIKE1, CYCLOIDEA, and PROLIFERATING CELL FACTOR1 | TCP (PF03634) | |
| ZF-HD | ZF-HD | Zinc finger homeodomain protein | ZF-HD_dimer (PF04770) | |
| SBP | SBP | SQUAMOSA promoter binding protein | SBP (PF03110) | |
| SRS | SRS | SHI RELATED SEQUENCE | DUF702 (PF05142) | |
| SAP | SAP | STERILE APETALA | SAP (PlantTFDB) | |
| Whirly | Whirly | Whirly | Whirly (PF08536) | |
| STAT | STAT | Signal Transducers and Activators of Transcription | STAT (PlantTFDB) | |
| VOZ | VOZ | Vascular plant One-Zinc finger | VOZ (PlantTFDB) | |

*DBD: DNA binding domain

Supplementary Table S3: Description of 59 TF genes identified by our pipeline and not found in PlantTFDB.

| Category | Gene Description | Genes |
|---|---|---|
| Genes annotated as transcription factors | C2H2 and C2HC zinc fingers superfamily protein | AT1G13400, AT1G68360, AT1G68480, AT3G01030, AT3G23140, AT4G04404, AT5G54340 |
| | B3 domain protein | AT1G50220, AT1G51970, AT1G78640, AT4G03160, AT4G05630, AT5G54067 |
| | Zinc finger protein 622 | AT2G24500, AT4G31420 |
| | C2H2-like zinc finger protein | AT5G48890, AT5G54360 |
| | Zinc finger C-x8-C-x5-C-x3-H type family protein | AT1G21580, AT5G66270 |
| | SET domain-containing protein | AT2G23380, AT4G02020 |
| | Homeodomain-like superfamily protein | AT2G21650, AT3G53440 |
| | response regulator 23 | AT5G62120 |
| | B3 DNA-binding domain protein | AT1G10455 |
| | bZIP family transcription factor | AT1G35490 |
| | Basic-leucine zipper (bZIP) transcription factor family protein | AT2G21235 |
| | bZIP protein | AT5G04840 |
| | C2H2-type zinc finger family protein | AT1G02040 |
| | C2H2 type zinc finger transcription factor family | AT1G49900 |
| | zinc finger protein 6 | AT1G67030 |
| | zinc finger (C2H2 type) family protein | AT2G36930 |
| | CCCH-type zinc finger family protein | AT2G02160 |
| | DNA-binding storekeeper protein transcriptional regulator- | AT2G20805 |
| | Homeodomain-like protein with RING/FYVE/PHD-type zinc | AT1G01150 |
| | ALWAYS EARLY 2 | AT5G27610 |
| | ADA2 2A | AT3G05380 |
| | ADA2 2B | AT3G07740 |
| | Protein ALWAYS EARLY 1 | AT4G16420 |
| | WRKY family transcription factor | AT3G32090 |
| Genes not annotated as transcription factors | F-box associated ubiquitination effector family protein | AT2G21920 |
| | Telomerase activator1 | AT3G09290 |
| | Protein arginine methyltransferase 3 | AT3G12270 |
| | Protein CTF7 | AT4G31400 |
| | D111/G-patch domain-containing protein | AT5G26610 |
| | Bromodomain 4 | AT1G61215 |
| | ELM2 domain-containing protein | AT2G03470 |
| | Histone H2A deubiquitinase (DUF3755) | AT3G07565 |
| | Duplicated homeodomain-like superfamily protein | AT3G12560 |
| | TRF-like 9 | AT3G46590 |
| | TRF-like 1 | AT4G11400 |
| | ARID/BRIGHT DNA-binding , ELM2 domain and myb-like | AT5G03780 |
| | TRF-like 10 | AT5G13820 |
| | Telomeric DNA binding protein 1 | AT5G59430 |
| | Telomeric repeat binding protein 1 | AT2G46280 |
| | TGF-beta receptor interacting protein 1 | AT2G46290 |
| | Transducin/WD40 repeat-like superfamily protein | AT5G01770 |
| | Regulatory-associated protein of TOR 2 (RAPTOR2) | AT5G51800 |
| | Protein kinase superfamily protein | AT2G21920 |

Supplementary Table S5: Twenty-eight orthologous groups from the AP2 family. OGs with variance equal or greater than 0.5 (marked with asterisks) were used for gene birth-death rate estimation in CAFE.

| **Species** | ***AP2:1** | ***AP2:2** | ***AP2:3** | ***AP2:4** | ***AP2:5** | ***AP2:6** | ***AP2:7** | ***AP2:8** | **AP2:9** | **AP2:10** | ***AP2:11** | ***AP2:12** | **AP2:13** | **AP2:14** | **AP2:15** | **AP2:16** | **AP2:17** | **AP2:18** | **AP2:19** | **AP2:20** | **AP2:21** | **AP2:22** | **AP2:23** | **AP2:24** | **AP2:25** | **AP2:26** | **AP2:27** | **AP2:28** |
|---|---|---|---|---|---|---|---|---|---|---|---|---|---|---|---|---|---|---|---|---|---|---|---|---|---|---|---|---|
| *Se. moellendorffii* | 0 | 0 | 0 | 0 | 0 | 0 | 0 | 0 | 0 | 0 | 0 | 1 | 5 | 0 | 3 | 0 | 0 | 0 | 0 | 0 | 0 | 0 | 0 | 0 | 0 | 0 | 0 | 2 |
| *Am. trichopoda* | 2 | 1 | 0 | 1 | 1 | 1 | 0 | 1 | 1 | 0 | 0 | 0 | 0 | 0 | 0 | 0 | 0 | 0 | 0 | 0 | 0 | 0 | 0 | 0 | 0 | 0 | 0 | 0 |
| *Aq. coerulea* | 3 | 2 | 1 | 2 | 2 | 2 | 1 | 1 | 1 | 0 | 0 | 0 | 0 | 0 | 0 | 0 | 0 | 0 | 0 | 0 | 0 | 0 | 0 | 0 | 0 | 0 | 0 | 0 |
| *Vi. vinifera* | 3 | 1 | 3 | 2 | 2 | 1 | 2 | 1 | 1 | 1 | 0 | 0 | 0 | 0 | 0 | 1 | 0 | 0 | 0 | 0 | 1 | 0 | 0 | 0 | 0 | 0 | 0 | 0 |
| *Ar. thaliana* | 4 | 1 | 1 | 2 | 0 | 2 | 2 | 1 | 1 | 1 | 0 | 0 | 0 | 0 | 0 | 0 | 0 | 0 | 0 | 0 | 0 | 2 | 0 | 0 | 0 | 0 | 0 | 0 |
| *Ch. fasciculata* | 3 | 1 | 3 | 2 | 3 | 2 | 2 | 0 | 2 | 1 | 1 | 0 | 0 | 0 | 0 | 0 | 0 | 0 | 0 | 0 | 0 | 0 | 0 | 0 | 0 | 0 | 0 | 0 |
| *Lo. japonicus* | 3 | 1 | 3 | 2 | 2 | 2 | 0 | 0 | 1 | 2 | 0 | 0 | 0 | 0 | 0 | 2 | 0 | 0 | 0 | 1 | 0 | 0 | 0 | 1 | 2 | 1 | 0 | 0 |
| *Gl. uralensis* | 4 | 4 | 3 | 2 | 3 | 2 | 1 | 2 | 0 | 1 | 1 | 0 | 0 | 0 | 0 | 0 | 0 | 0 | 0 | 0 | 0 | 0 | 0 | 0 | 0 | 1 | 0 | 0 |
| *Me. truncatula* | 4 | 4 | 3 | 3 | 3 | 2 | 2 | 2 | 1 | 1 | 1 | 0 | 0 | 0 | 0 | 0 | 0 | 0 | 0 | 0 | 0 | 0 | 0 | 0 | 0 | 0 | 2 | 0 |
| *Ci. reticulatum* | 4 | 4 | 3 | 1 | 3 | 2 | 2 | 1 | 1 | 1 | 1 | 0 | 0 | 0 | 0 | 0 | 0 | 0 | 0 | 0 | 0 | 0 | 1 | 0 | 0 | 0 | 0 | 0 |
| *Ci. arietinum* | 4 | 4 | 3 | 2 | 3 | 2 | 2 | 1 | 1 | 1 | 1 | 0 | 0 | 0 | 0 | 0 | 0 | 0 | 0 | 0 | 0 | 0 | 1 | 0 | 0 | 0 | 0 | 0 |
| *Ca. cajan* | 4 | 4 | 3 | 4 | 2 | 2 | 2 | 1 | 1 | 1 | 1 | 0 | 0 | 0 | 0 | 0 | 0 | 0 | 0 | 0 | 0 | 0 | 0 | 0 | 0 | 0 | 0 | 0 |
| *Gl. soja* | 7 | 8 | 6 | 6 | 4 | 3 | 4 | 2 | 2 | 2 | 2 | 1 | 0 | 1 | 0 | 0 | 1 | 1 | 0 | 0 | 0 | 0 | 0 | 1 | 0 | 0 | 0 | 0 |
| *Gl. max* | 6 | 8 | 6 | 5 | 3 | 4 | 4 | 2 | 3 | 2 | 2 | 1 | 0 | 1 | 0 | 0 | 1 | 1 | 0 | 0 | 0 | 0 | 0 | 0 | 0 | 0 | 0 | 0 |
| *Ph. vulgaris* | 3 | 4 | 3 | 4 | 2 | 3 | 2 | 1 | 1 | 1 | 1 | 1 | 0 | 1 | 0 | 0 | 1 | 1 | 0 | 0 | 0 | 0 | 0 | 0 | 0 | 0 | 0 | 0 |
| *Vi. radiata* | 3 | 4 | 3 | 4 | 2 | 2 | 2 | 1 | 1 | 1 | 1 | 0 | 0 | 0 | 0 | 0 | 0 | 0 | 0 | 0 | 0 | 0 | 0 | 0 | 0 | 0 | 0 | 0 |
| *Vi. angularis* | 3 | 4 | 3 | 4 | 2 | 2 | 2 | 1 | 1 | 1 | 1 | 1 | 0 | 1 | 0 | 0 | 0 | 0 | 0 | 0 | 0 | 0 | 0 | 0 | 0 | 0 | 0 | 0 |
| *Lu. angustifolius* | 9 | 4 | 5 | 2 | 0 | 4 | 2 | 3 | 2 | 3 | 3 | 0 | 0 | 0 | 0 | 0 | 0 | 0 | 0 | 0 | 0 | 0 | 0 | 0 | 0 | 0 | 0 | 0 |
| *Ar. duranensis* | 5 | 4 | 3 | 2 | 3 | 2 | 2 | 1 | 1 | 1 | 0 | 0 | 0 | 0 | 0 | 0 | 0 | 0 | 0 | 0 | 0 | 0 | 0 | 0 | 0 | 0 | 0 | 0 |
| *Ar. ipaensis* | 5 | 3 | 3 | 2 | 3 | 2 | 3 | 2 | 1 | 1 | 0 | 0 | 0 | 0 | 0 | 0 | 0 | 0 | 3 | 1 | 1 | 0 | 0 | 0 | 0 | 0 | 0 | 0 |
